# Characterization of bacterial fucokinase/GDP-fucose pyrophosphorylase (FKP) enzymes supports the evolution of interdomain communication and modularity in the FKP family

**DOI:** 10.1101/2025.08.05.668714

**Authors:** Nicholas T. Smith, Autumn J. Hodgins, Marie E. Boddington, Chantelle J. Capicciotti, George C. diCenzo, Graeme W. Howe

## Abstract

L-fucokinase (FUK) and GDP-fucose pyrophosphorylase (GFPP) salvage free L-fucose and synthesize the valuable nucleotide-sugar GDP-L-fucose (GDP-Fuc). Some organisms express these enzymes as one bifunctional polypeptide called L-fucokinase/GDP-fucose pyrophosphorylase (FKP), which has attracted attention for use in the chemoenzymatic synthesis of GDP-Fuc. Despite the documented use of the FKP from *Bacteroides fragilis* (*Bf*FKP), the evolutionary origins of these enzymes and their relationships to monofunctional FUKs and GFPPs are poorly understood. We hypothesized that biochemical characterization of these proteins coupled with an evolutionary analysis would uncover the natural diversity of FKPs, facilitating the discovery of new biocatalysts. Phylogenetic and sequence similarity network (SSN) analyses distinguished FKPs from FUKs and GFPPs, suggesting that FKPs originate from one ancestral fusion event between these domains. To evaluate how environmental factors might select for functional diversity within the FKP family, we recombinantly expressed and purified a putative FKP from the thermophilic bacterium *Thermophagus xiamenensis* (*Tx*FKP). This enzyme exhibited *in vitro* kinase and pyrophosphorylase activities and demonstrated subtle kinetic differences compared to *Bf*FKP. While alanine scanning mutational analysis of the *Tx*FKP FUK and GFPP domains supported the role of conserved residues that *Tx*FKP uses to coordinate substrate binding and catalysis, other mutations in the *Tx*FKP GFPP domain influenced kinase activity differentially for the substrates L-fucose and D-arabinose, showing an unprecedented role for the GFPP domain in FUK substrate specificity. Finally, thermal shift profiles of *Tx*FKP and *Bf*FKP were biphasic and provided new insights into how these enzymes have evolved to respond to different sugar substrates.

## INTRODUCTION

Fucosylation plays a vital role in processes such as cancer progression (1), pathogenesis (2), and plant-microbe signalling (3). Given the myriad roles of L-fucose (L-Fuc), the enzymes responsible for the installation of this sugar (fucosyltransferases) have received significant attention. Although these proteins have evolved diverse acceptor specificities, all known fucosyltransferases employ the nucleotide sugar guanosine 5′-diphosphate-β-L-fucose (GDP-Fuc) as the donor substrate (4). Therefore, the metabolism of GDP-Fuc is intertwined with the outcome of fucosylation, or dysfunction thereof, in broad biological scenarios. GDP-Fuc is also prohibitively expensive (5), presenting a barrier for *in vitro* fucosyltransferase research and necessitating an improved understanding of the enzymes responsible for GDP-Fuc production (5–7).

GDP-Fuc is biosynthesized naturally via an evolutionarily conserved *de novo* pathway, in which the enzymes GDP-D-Man-4,6-dehydratase and GDP-4-keto-6-deoxy-D-Man-3,5-epimerase-4- reductase convert GDP-D-Mannose to GDP-Fuc over three steps (8–10). However, GDP-Fuc is alternatively synthesized through a salvage pathway that uses fucokinase (FUK) and GDP-fucose pyrophosphorylase (GFPP) to “salvage” free fucose into GDP-Fuc (11–13). In this pathway, FUK phosphorylates L-Fuc using ATP, and the resulting fucose-1-phosphate (Fuc-1-P) is converted to GDP-Fuc by the GFPP enzyme. The salvage pathway only accounts for approximately 10% of GDP-Fuc production in mammals (14), yet cells can seemingly discriminate between GDP-Fuc synthesized by these different pathways, and sometimes prefer salvaged pools of GDP-Fuc over GDP-Fuc obtained from *de novo* synthesis (15, 16). Both differential regulation and perturbation of the salvage pathway have clinical implications in humans (17, 18), suggesting that this alternative biosynthetic route can play important, context-specific roles. The fucose salvage pathway was thought to be restricted to eukaryotes until Coyne et al. reported an analogous bacterial pathway in 2005 (19). The salvage pathway has now been reported across eukaryotic and prokaryotic taxa including fungi (20, 21), plants (22, 23), protists (24), and bacteria (19, 25), making it an adaptation widely used in Nature.

Intriguingly, some organisms express both components of the salvage pathway as a single, bifunctional enzyme called L-fucokinase/GDP-fucose pyrophosphorylase (FKP) (19). This enzyme was first discovered in the mammalian gut symbiont *Bacteroides fragilis* 9343 (*Bf*FKP), by virtue of sequence similarity of the *N*- and *C*-termini to known mammalian GFPPs and FUKs, respectively (19). As FKP catalyzes both steps in the salvage pathway, this protein has emerged as an attractive tool for affordable GDP-Fuc production (6), with recent studies focussing on improving the chemoenzymatic synthesis of GDP-Fuc by FKP, allowing for increased yields and the synthesis of sugars with unnatural substituents (5, 26). Conditions for gram-scale production of GDP-Fuc using *Bf*FKP and continuous recycling of ATP have been described (5), reflecting the importance of access to this metabolite for the study of fucosylation and fucosyltransferase enzymes responsible for the installation of this critical sugar. Further, heterologous expression of *Bf*FKP in the yeast *Saccharomyces cerevisiae* imbued transformants with the ability to produce GDP-Fuc, demonstrating that FKP enzymes can fulfill the salvage pathway role when engineered into unnatural hosts (27). Since its identification, *Bf*FKP has been kinetically characterized (6, 26, 28, 29), residues important for substrate binding and catalysis have been elucidated (28, 29), and the quaternary structure has been determined (28–30). *Bf*FKP has also been found to exhibit a degree of promiscuity with the acceptor sugar substrates, facilitating the chemoenzymatic production of GDP-sugars with various C5-substituted L-Fuc analogs, including D-arabinose (D-Ara) (26).

Despite the importance of FKP for the production of GDP-Fuc, *Bf*FKP is the sole bacterial FKP protein that has been characterized, and studies on FKP orthologs from other organisms are scarce. The *At*FKGP protein from the model plant *Arabidopsis thaliana* also catalyzes the synthesis of GDP-Fuc from L-Fuc, but the FUK domain of this homolog possesses a higher affinity for ATP and a significantly lower affinity for L-Fuc than *Bf*FKP (22). The enzymes FKP40 and AFKP80 from the protozoan parasite *Leishmania major* also salvage L-Fuc, although AFKP80 is capable of fully converting D-Ara to GDP-D-arabinose (GDP-Ara), while FKP40 only phosphorylates D-Ara and displays almost no pyrophosphorylase activity necessary for installation of the GDP moiety (24, 31).

Although FKP proteins have proven useful for both biotechnological use and fundamental glycobiology, the evolutionary history of this protein family has not been explored, and the abundance of FKP orthologs compared to monofunctional FUKs and GFPPs is unclear. Since *B. fragilis* fucosylates its surface polysaccharides and cannot persist within the gut microbiome without GDP-Fuc, it was first hypothesized that this organism’s FKP provides it with a competitive advantage in the gut environment (19). However, the presence of FKP enzymes in organisms from other environments suggests that these enzymes may also provide fitness benefits absent from the context of the gut microbiota. To address these questions, we conducted a thorough bioinformatic analysis of the FKP, FUK, and GFPP protein families. A combination of phylogenetic inference and sequence similarity networks (SSNs) revealed multiple underexplored families of FKP proteins that likely originated from a single fusion event. We show that a putative FKP protein from the bacterium *Thermophagus xiamenensis* (dubbed *Tx*FKP) is an authentic FKP, representing only the second characterized bacterial ortholog. *In vitro* kinetic characterization and thermal shift profiling of *Bf*FKP and *Tx*FKP were used to demonstrate potential interdomain regulation patterns and differences in conformational responses to substrate binding. These results show that while many *Tx*FKP properties were conserved with *Bf*FKP, interdomain communication and conformational dynamics may have evolved differently following the fusion of their monofunctional ancestors.

## RESULTS AND DISCUSSION

### Evolution and diversity of FKP proteins

To better understand the evolutionary history of FKP proteins, we queried the >63 million proteins of the UniRef50 database (i.e., representative proteins of UniRef90 proteins clustered at 50% sequence identity, with one protein per cluster) for entries that matched Hidden Markov Models (HMMs) for FUKs, GFPPs, and bifunctional FKP proteins. Testing the hypothesis that FKP proteins originated from a fusion between FUKs and GFPPs, we first analyzed these domains independently. We generated separate phylogenies with the FUK and GFPP proteins and used IQ-TREE2’s non-reversible model to estimate the root positions (32). The resulting FUK phylogeny had a monophyletic group of FKP proteins that were distinct from most of their monofunctional counterparts (Figure 1A), while the estimated root in the GFPP phylogeny was ambiguous. However, the GFPP phylogeny could be rooted such that all FKP proteins were monophyletic (Figure 1B), or at its midpoint, resulting in the FKP proteins being distributed across three separate clades (Figure S1). Since it is more parsimonious that the sole ancestral FUK fused with one ancestral GFPP rather than multiple separate GFPPs, we favour the hypothesis that all known FKPs evolved from a single fusion event.

**Figure 1.**
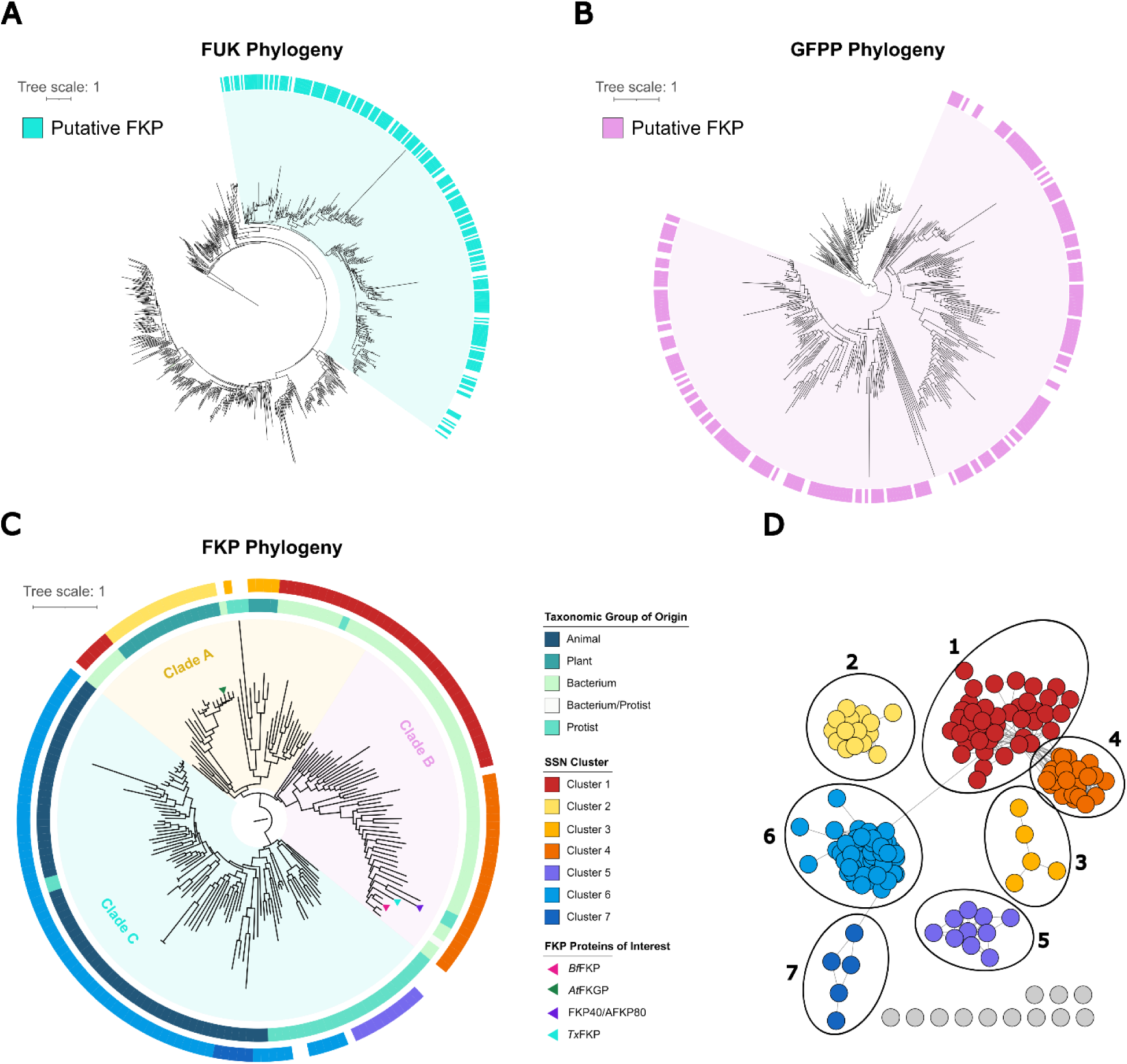
Evolution of bifunctional FKP proteins. A combination of bioinformatic analyses demonstrating relationships between bifunctional FKP proteins, FUKs, and GFPPs are shown. For all phylogenies the scale bar represents the average number of amino acid substitutions per site, node support was calculated based on 1000 ultrafast bootstrap replicates, and interactive forms with bootstrap supports can be accessed at https://itol.embl.de/shared/1tzSBFA310S3y. (A) A maximum-likelihood phylogeny based on UniRef50 hits to the FUK protein family. Protein clusters that also have predicted GFPP domains, and thus are putative FKP proteins, are indicated with a cyan rectangle. The “FKP monophyletic clade” is highlighted in light cyan. (B) A maximum-likelihood phylogeny generated from UniRef50 hits to the GFPP protein family. Protein clusters that also have predicted FUK domains, and thus are putative FKP proteins, are indicated with a pink rectangle. The “FKP monophyletic clade” is highlighted in light pink. (C) A maximum-likelihood phylogeny generated from predicted bifunctional FKP UniRef50 clusters. Only accessions represented by sequences between 745 and 1745 amino acids in length were included. Major clades of interest (A, B, or C) are indicated using yellow (Clade A), pink (Clade B), or blue (Clade C) shading. Data strips (coloured rectangles) indicate sequence similarity network clusters or taxonomic groupings. Proteins that do not cluster in the sequence similarity network are uncoloured. UniRef50 accessions containing FKP proteins of interest are shown with arrows, as indicated in the legend. (D) A sequence similarity network generated from the same FKP protein set that was used for panel C. Nodes (circles) represent UniRef50 accessions, and the connecting edges (lines) represent a shared identity of approximately 30% or above. Distinct clusters are annotated and coloured accordingly.

Interestingly, some monofunctional FUKs and GFPPs grouped with the FKP clade in both phylogenies and the accompanying SSNs (Figure 1A, 1B, Figure S2). This indicates that these proteins were, surprisingly, more closely related to the FKP proteins than to most other FUKs or GFPPs. For example, the characterized FUK and GFPP proteins from the fungus *Mortierella alpina* (21, 33) were found in the FKP monophyletic clade in both the FUK and GFPP phylogenies, suggesting that both have common ancestry with the FKP proteins, despite no fungal FKP enzymes being detected by our analyses. These *M*. *alpina* proteins are also thought to share common ancestry with the *Amoebozoa* salvage pathway proteins (20), a phylum in which FKP enzymes were detected. Therefore, our data suggest that a subset of extant FUKs and GFPPs are descendants of FKP enzymes. The data thus illuminate a complex evolutionary history in which FKP enzymes arose from one fusion event and are modified, lost, and/or differentially regulated in some organisms.

If all known FKP proteins originated from a single fusion event, their functional diversity could be shaped partly by selective pressure. It was therefore of interest to model the evolution of bifunctional FKP proteins to source FKP proteins with potential functional divergence. A phylogeny generated from the putative FKP proteins contained three major clades, annotated as A, B, and C (Figure 1C). An SSN built from the same set of proteins was congruent with the phylogeny and revealed groupings specific to certain organisms (Figure 1C, 1D). Clade A contained FKPs from plants (such as *At*FKGP), bacteria, and a few protists. Clusters 2 and 3, which include all identified plant FKPs, were nested within the clade of bacterial proteins from Cluster 1 in the phylogeny, suggesting that the corresponding genes have been acquired horizontally from bacteria. The plant proteins in Cluster 3 originate only from chlorophytic algae, while Cluster 2 plant proteins originate from charophytic algae and embryophyte land plants like mosses and vascular plants. As chlorophytes likely diverged from the plant lineage before charophytes (34), our data supports separate FKP acquisitions that occurred after this divergence. Interestingly, the *At*FKGP protein was originally hypothesized to share ancestry with bacterial FKP proteins (22). Our analysis agrees with this hypothesis and further suggests that the plant FKP proteins were acquired horizontally from bacteria.

Clade B was saturated with bacterial proteins, many of which were from the phylum *Bacteroidota* (including *Bf*FKP), reflecting the initial observation that FKP proteins are prevalent in microbes commonly associated with the gut (19). The characterized FKP40/AFKP80 proteins from *L. major* and a putative FKP protein from the protist *Strigomonas culicis* are also in Clade B, where they are close relatives to *Bf*FKP. Further, several FKP proteins from *Trypanosoma cruzi* are members of the same UniRef50 group as *Bf*FKP. *L*. *major*, *S*. *culicis*, and *T*. *cruzi* belong to the *Trypanosomatida* order of protozoan parasites, representing the only non-bacterial proteins in Clade B. This suggests another horizontal transfer of FKP genes from bacteria to eukaryotes, and agrees with previous reports of horizontal transfers of sugar kinase genes from bacteria to *Trypanosomatida* parasites (35).

The proteins within Clade C originated only from eukaryotes, and the majority of these were from animals. The proteins in Clusters 5 and 7 specifically originated from diatoms and nematodes, respectively. As salvage pathway enzymes are easily lost when they do not provide a fitness benefit (20, 35), it is possible that the clusters of FKP sequences unique to certain taxa reflect an important role for fucose salvage in these organisms. However, more evidence would be required for the careful evaluation of this hypothesis.

Overall, the phylogenetic analyses indicate that FKP proteins are abundant in some organisms, have been acquired horizontally by others, and that functional differences between FKP proteins may be the result of selective pressure. To complement this evolutionary analysis, we chose a putative FKP from the bacterium *Thermophagus xiamenensis* (*Tx*FKP; Uniprot accession ID: A0A1I2DJ10) for *in vitro* characterization. We were particularly interested in this protein because it is closely related to FKPs from gut-bacteria and intestinal parasites, but was sourced from a thermophilic organism that lives in hot spring sediments (36). Thus, *Tx*FKP was expected to exhibit enhanced thermostability and/or altered kinetic properties due to the ecological niche of its host organism. *Tx*FKP and *Bf*FKP share 51% sequence identity, and ColabFold2-generated models of the two proteins aligned with an RMSD of 1.154 Å across 786 pruned atom pairs (of 946 total atom pairs) (Figure 2A). Predicted substrate binding residues (Figure 2B, 2C) (28) were conserved in sequence and structural models across both enzymatic domains of the proteins (Figure 2, Figure S3), providing *in silico* evidence that *Tx*FKP is likely an authentic L-fucokinase/GDP-Fucose pyrophosphorylase.

**Figure 2.**
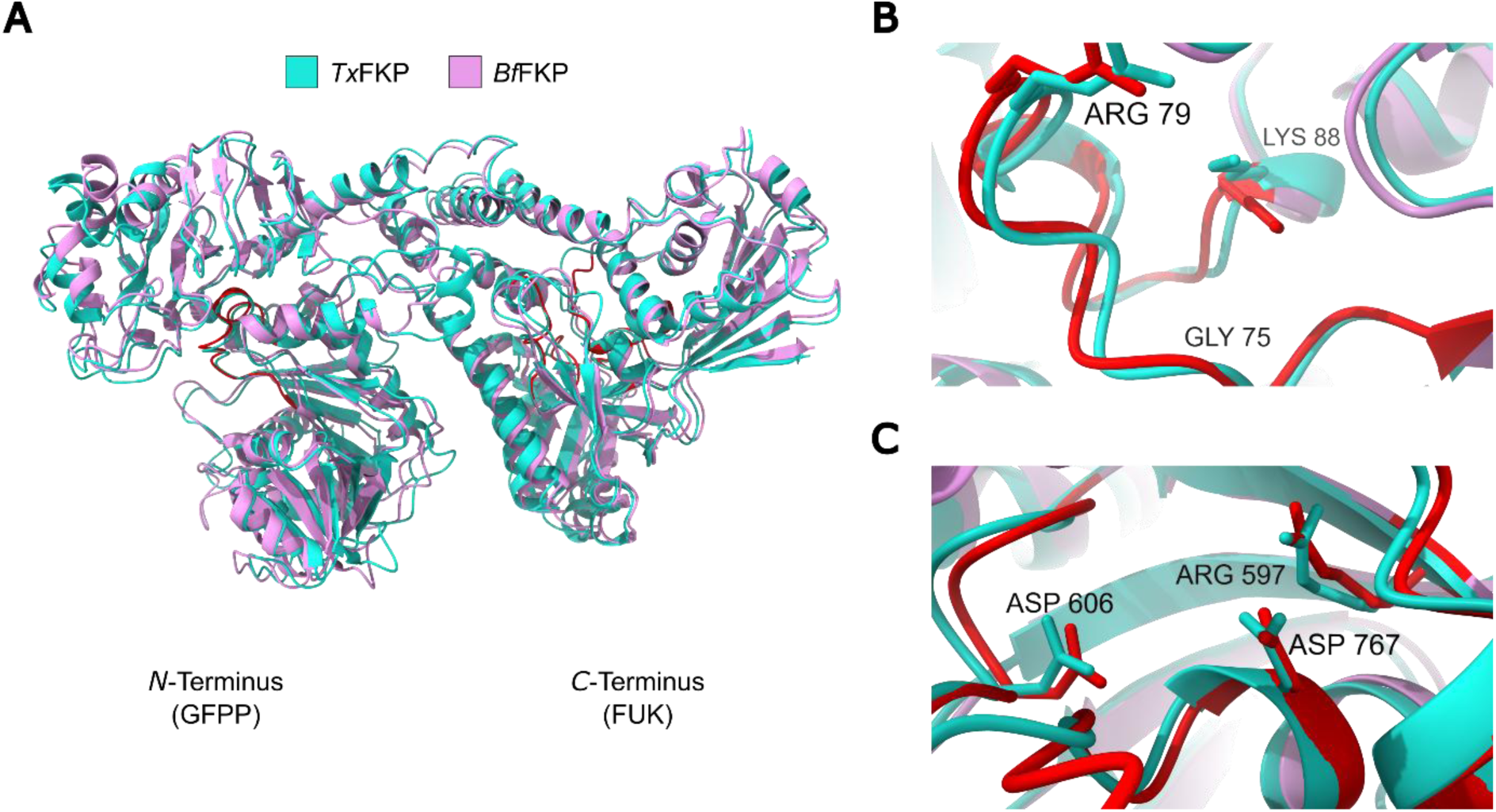
Structural conservation between ***Bf***FKP and ***Tx***FKP. A comparison of *Bf*FKP (pink) and *Tx*FKP (cyan) ColabFold2 (57) models. *Bf*FKP residues coloured in red are putative substrate binding residues identified by Liu et al (2019) (28). (A) Full-length structural alignment, performed with ChimeraX matchmaker (58). (B) Snapshot of the aligned N-terminal pyrophosphorylase domains. *Bf*FKP residues coloured in red are putative GTP-binding residues. Three residues that are critical for GFPP function in *Bf*FKP, and which were mutated in *Tx*FKP in this study, are labelled. Amino acid side- chains are only shown for residues targeted for mutagenesis. (C) Snapshot of the aligned C-terminal FUK domains. *Bf*FKP residues coloured in red are putative ATP or fucose-binding residues. Three residues that are critical for FUK function in *Bf*FKP, and which were mutated in *Tx*FKP in this study, are labelled. Amino acid side-chains are only shown for residues targeted for mutagenesis.

### Recombinant expression and purification of BfFKP and TxFKP

Throughout our analysis of *Tx*FKP, we incorporated a *Bf*FKP control to provide a comparison under identical experimental conditions. Both FKP proteins were recombinantly expressed and purified with immobilized metal affinity chromatography (IMAC) (Table S4, Figure S4). Co-expression with the chaperone proteins GroEL, GroES, and Trigger Factor (TF) greatly increased the yield and purity of soluble *Tx*FKP. Attempts to overexpress *Tx*FKP without co-expression of these chaperones were unsuccessful, while repeated trials to purify *Tx*FKP from co-expression cultures were contaminated with a ∼60 kDa protein that likely corresponds to the GroEL chaperone. A similar co-purification issue with GroEL was reported in the recombinant purification of *At*FKGP (22). Despite literature precedent (29), attempts to separate the contaminant from *Tx*FKP by incubating the mixture with denatured proteins and ATP were unsuccessful. Nonetheless, the usage of chaperone proteins circumvented the need to generate FKP fusions with solubility tags, a strategy that has been used for *Bf*FKP and is predicted to alter the kinetic activity or oligomerization of these fusions (6, 29). To maintain consistency, we co-expressed both *Tx*FKP and *Bf*FKP in the presence of the chaperones described above, although little co-purification was observed with *Bf*FKP.

### TxFKP-catalyzed GDP-fucose synthesis

To determine if *Tx*FKP can convert L-Fuc into GDP-Fuc, *Tx*FKP was incubated with L-Fuc, ATP, GTP, MgCl2, and MnCl2 at 37 °C for one hour. After quenching the reactions, the mixtures were analyzed by HILIC-HPLC-MS (37, 38) to characterize the products formed. Initial experiments using *Bf*FKP and L-Fuc as the substrate confirmed that this method could accurately detect Fuc-1-P and GDP-Fuc products, which eluted at retention times of approximately 28.5 and 29.6 minutes, respectively (Figure S5, S6). The ability of *Tx*FKP to convert L-Fuc into its products in a multi-step reaction was next assessed (Figure 3). In a negative control reaction, unmodified L-Fuc was detected with a retention time of approximately 10 minutes, whereas only residual L-Fuc could be observed in the experimental reaction. In the *Tx*FKP reaction, GDP-Fuc was clearly detected, while Fuc-1-P was not. This suggested that any produced Fuc-1-P had been rapidly consumed and converted to GDP-Fuc under the assay conditions (Figure 3, Figure S8). These results indicate that the FUK and GFPP domains were both active and were capable of acting together to convert L-Fuc into GDP-Fuc. *Tx*FKP was capable of producing GDP-Fuc in the presence of 10 mM MgCl2, 10 mM MnCl2, or 10 mM of both salts, showing that both domains can use either Mg^2+^ or Mn^2+^ cations for their respective reactions. In reactions performed in the absence of either metal and in the presence of 5 mM EDTA, neither Fuc-1-P nor GDP-Fuc were detected. We also confirmed the previous report (26) that this requirement was shared by *Bf*FKP (Figure S5, S6), establishing the absolute requirement for a divalent metal cation as a common feature of these FKP enzymes.

**Figure 3.**
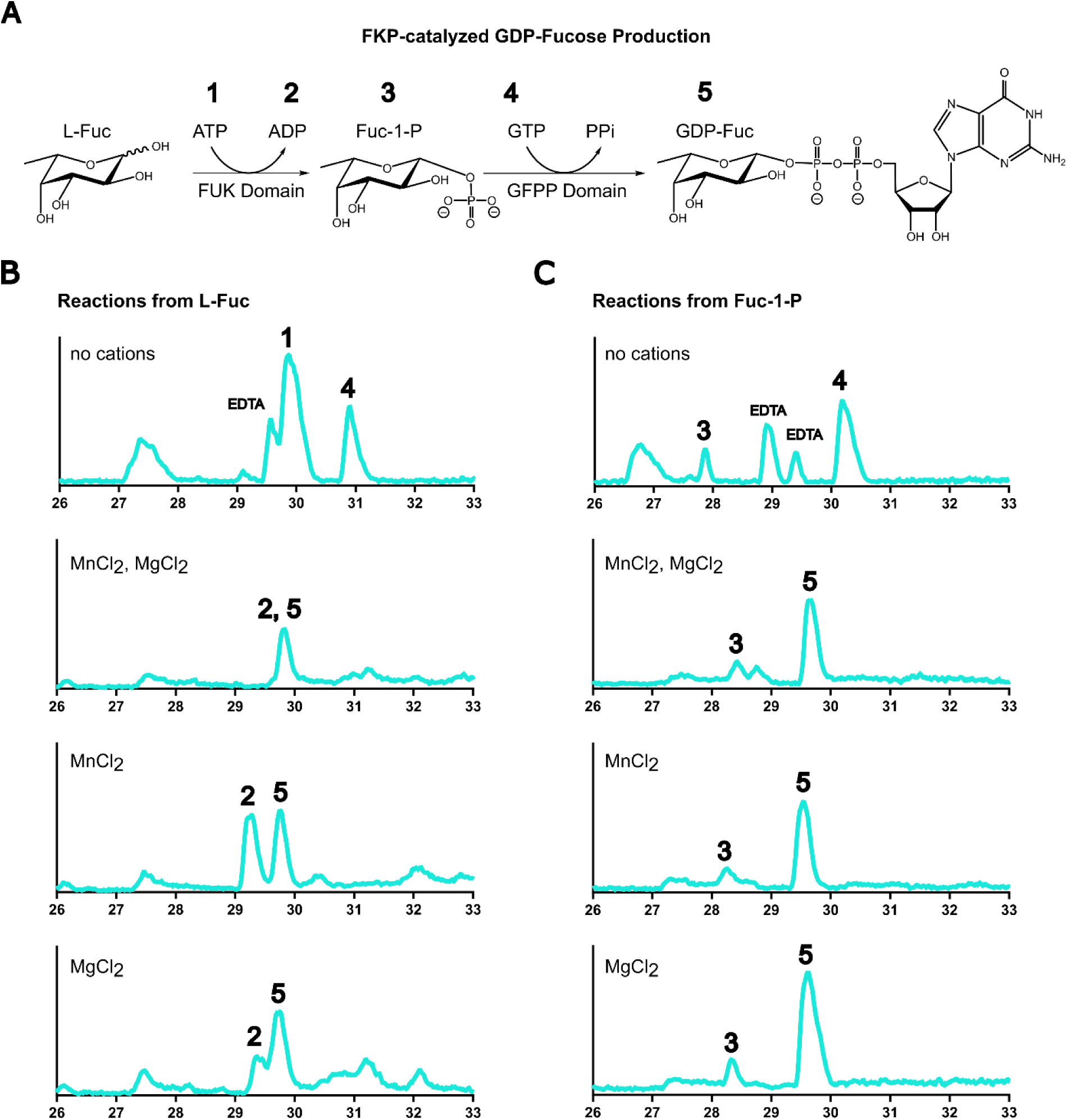
***Tx***FKP-catalyzed GDP-fucose synthesis. HILIC-HPLC results supporting the *Tx*FKP-catalyzed synthesis of GDP-Fuc are shown. Numbers from one to five represent a reaction substrate or product as follows: (1) ATP, (2) ADP, (3) Fuc-1-P, (4) GTP, and (5) GDP-Fuc. Mass spectrometry chromatograms (negative mode), with signal intensity plotted against time, are shown of reaction mixtures following one hour of incubation at 37 °C. All chromatograms were standardized based on the retention time of HEPES (23-25 min). The salt(s) used in each condition are indicated on the chromatograms and the “no cation” condition included 5 mM EDTA. Relevant peaks are numbered accordingly and the peaks corresponding to EDTA is labelled where applicable. (A) A reaction scheme representing the FUK and GFPP-catalyzed reactions during the two-step synthesis of GDP-Fuc. (B) Chromatograms of reaction mixtures where *Tx*FKP was incubated with L-Fuc as a starting substrate. (C) Chromatograms of reaction mixtures where *Tx*FKP was incubated with Fuc-1-P as a starting substrate.

We also asked if *Tx*FKP could produce GDP-Fuc using exogenously provided Fuc-1-P as substrate. To probe this possibility, the enzyme was incubated with Fuc-1-P, GTP, MgCl2, and MnCl2 at 37 °C for one hour. Again, GDP-Fuc was detected in both the *Tx*FKP and *Bf*FKP reactions (Figure 3C, Figure S5, S7, S9), showing that *Tx*FKP can bind and process exogenously supplied Fuc-1-P into GDP-Fuc. The *Tx*FKP-catalyzed conversion of Fuc-1-P into GDP-Fuc occurred in the presence of 10 mM Mg^2+^ and/or 10 mM Mn^2+^, but not in the absence of these cations, supporting our initial observation that the *Tx*FKP pyrophosphorylase activity is cation-dependent. Collectively, these results demonstrate that *Tx*FKP is an authentic FKP enzyme that can synthesize GDP-Fuc from L-Fuc (i.e., the fucose salvage pathway), or from the intermediate Fuc-1-P, *in vitro*.

### Kinetic analysis of the TxFKP-catalyzed reactions

To monitor the kinetics of the *Tx*FKP-catalyzed reactions, we employed two coupled assays with spectrophotometric readouts. The fucokinase reaction was monitored using a coupled assay that used phosphoenolpyruvate (PEP), NADH, pyruvate kinase (PK) and lactate dehydrogenase (LDH). This coupled assay allows for initial rates of ADP production (and thus Fuc-1-P production; Figure 3A) to be measured using absorbance decreases at 340 nm (39). The pyrophosphorylase reaction was performed in the presence of the inorganic pyrophosphatase from *Pasteurella multocida* (*Pm*PpA) (40), enabling the breakdown of inorganic pyrophosphate (PPi) into inorganic phosphate. The reactions were incubated with malachite green detection reagent and quantified with an increase in absorbance at 635 nm (41).

We first examined the affinity of the *Tx*FKP *C*-terminal kinase domain for its substrates. Since FKP enzymes from Clade B in our evolutionary model are known to accept C5-modified L-Fuc derivates (24, 26), we tested activity on the biologically relevant sugars L-Fuc and D-Ara, which differ only by the presence of a C5 methyl group. Table 1 summarizes the kinetic parameters measured for the FKP-catalyzed kinase reactions. *Tx*FKP exhibited similar *K*M values for L-Fuc and D-Ara as *Bf*FKP, which were in agreement with previous characterizations of *Bf*FKP (26, 28) (Table 1, Figure S10). *Tx*FKP had a much higher affinity for L-Fuc than D-Ara, and the magnitude of the second order rate constants (*k*cat*/K*M) obtained with L-Fuc and D-Ara suggest that fucose phosphorylation is more likely its biological role, while arabinose phosphorylation is likely a promiscuous activity. However, *Tx*FKP catalyzed the phosphorylation of D-Ara with a *k*cat of nearly double that of the phosphorylation of L-Fuc, a feature we did not observe with *Bf*FKP. This difference might suggest that *Tx*FKP does have some *in vivo* role for D-Ara phosphorylation and has evolved a higher turnover rate to counteract its low affinity for this substrate. This would not be the only case where FKP enzymes have been co-opted for use with D- Ara; *L*. *major* depends on its AFKP80 protein to produce GDP-Ara *in vivo* (24). However, a more comprehensive analysis of Clade B FKPs would be required to test the conservation of this property.

**Table 1.** Kinetic parameters of *Tx*FKP and *Bf*FKP.

| | $K_M$ ($\mu\text{M}$) | $k_{\text{cat}}$ ($\text{s}^{-1}$) | $k_{\text{cat}}/K_M$ ($\text{M}^{-1} \text{s}^{-1}$) |
| --- | --- | --- | --- |
| ***Tx*FKP Kinase Activity** |  |  |  |
| L-fucose | $77.30 \pm 18.59$ | $0.90 \pm 0.07$ | $(1.2 \pm 0.3) \times 10^4$ |
| D-arabinose | $1322.00 \pm 150.10$ | $1.33 \pm 0.07$ | $(1.0 \pm 0.1) \times 10^3$ |
| ATP | $1035.00 \pm 348.00$ | $0.74 \pm 0.07$ | $(7.2 \pm 2.5) \times 10^2$ |
| ***Bf*FKP Kinase Activity** |  |  |  |
| L-fucose | $90.61 \pm 14.99$ | $0.94 \pm 0.06$ | $(1.0 \pm 0.2) \times 10^4$ |
| D-arabinose | $1783.00 \pm 258.00$ | $0.74 \pm 0.04$ | $(4.1 \pm 0.6) \times 10^2$ |
| ATP<sup>†</sup> | N.D. | N.D. | N.D. |
| ***Tx*FKP Pyrophosphorylase Activity** |  |  |  |
| Fucose-1-phosphate | $66.35 \pm 21.53$ | $4.81 \pm 0.62$ | $(7.3 \pm 2.5) \times 10^4$ |
| GTP | $26.85 \pm 11.20$ | $8.04 \pm 1.07$ | $(3.0 \pm 1.3) \times 10^5$ |
| ***Bf*FKP Pyrophosphorylase Activity** |  |  |  |
| Fucose-1-phosphate | $115.90 \pm 60.94$ | $8.63 \pm 2.71$ | $(7.4 \pm 4.6) \times 10^4$ |
| GTP | $138.70 \pm 31.93$ | $12.93 \pm 3.09$ | $(9.3 \pm 3.1) \times 10^4$ |

<sup>†</sup>Parameters not determined due to the observed inhibitory effect.

While characterizing the kinase activity of *Bf*FKP, we found that concentrations of ATP greater than 3 mM suppressed the rate of the *Bf*FKP-catalyzed fucokinase reaction (Figure S11). This apparent substrate inhibition pattern had not been previously reported for *Bf*FKP. With [ATP] held at 5 mM, the concentration of *Bf*FKP was proportional to the initial rate of reaction (Figure S12), indicating that the observed inhibition was intrinsic to *Bf*FKP and not caused by inhibition of the coupling enzymes. It is unclear why this inhibitory behaviour had not been reported before (26, 28), but previous kinetic characterizations of this enzyme may not have used a sufficiently wide array of [ATP] relative to the concentration of *Bf*FKP to reveal this facet of *Bf*FKP catalysis. While this phenomenon warrants further study, an investigation of *Bf*FKP inhibition is beyond the scope of this work. For our purposes, we opted to measure kinetic parameters for the *Bf*FKP-catalyzed phosphorylation of L-Fuc and D-Ara at a constant [ATP] = 3 mM, the concentration at which the maximum initial rate was observed (Table 1).

In contrast to *Bf*FKP, *Tx*FKP exhibited typical Michaelis-Menten kinetics up to >5x the apparent *K*M for ATP (Figure S10). However, much higher [ATP] did eventually reduce the initial rates of the *Tx*FKP- catalyzed reactions (Figure S13), suggesting that some form of substrate inhibition may be a conserved regulatory mechanism for FKP enzymes, at least those within Clade B.

Next, pyrophosphorylase kinetic assays of the *N*-terminal pyrophosphorylase domain revealed that *Tx*FKP had lower apparent *K*M values for Fuc-1-P and GTP than did *Bf*FKP; however, the values obtained with *Bf*FKP are higher than most previous reports suggest. Nonetheless, since the FKPs were evaluated under identical assay conditions, these results suggest that *Tx*FKP has a higher apparent affinity for its pyrophosphorylase substrates than *Bf*FKP does. *Tx*FKP also had a significantly higher *k*cat*/K*M value for the pyrophosphorylase activity than for the kinase activity (Table 1), suggesting that phosphorylation may be the rate-limiting step in the *Tx*FKP-catalyzed GDP-fucose salvage. This is consistent with the *k*cat*/K*M value we obtained with *Bf*FKP, and with the previous analysis of *At*FKGP (22), potentially highlighting another conserved feature between FKP enzymes from various clades in our phylogeny.

### The role of conserved TxFKP residues

Next, we evaluated the roles of key residues that are conserved between *Tx*FKP and other FKP enzymes. The residues targeted were previously reported to strongly influence either kinase or pyrophosphorylase activity in *Bf*FKP (28), so we rationalized that these amino acids would also be important for *Tx*FKP activity. We generated point mutations in both the *Tx*FKP *N*-terminus (K88A, G75A, R79A) and *C*-terminus (R597A, D606A, D767A) (Table S2, S4, Figure S4). All mutants expressed at levels similar to the wildtype except for K88A (Table S4), which did not express in the soluble form at a sufficient level for purification, potentially reflecting the role of the N-terminus in the solubility of FKP proteins (29). The activities of the mutant proteins were quantified using the same kinetic assays described above. Broadly, mutations in the *N*-terminal domain were deleterious to pyrophosphorylase activity and mutations in the *C*-terminal domain were deleterious to kinase activity (Figure 4), confirming that the *N*- and *C*-termini of *Tx*FKP confer distinct pyrophosphorylase and kinase activities.

**Figure 4.**
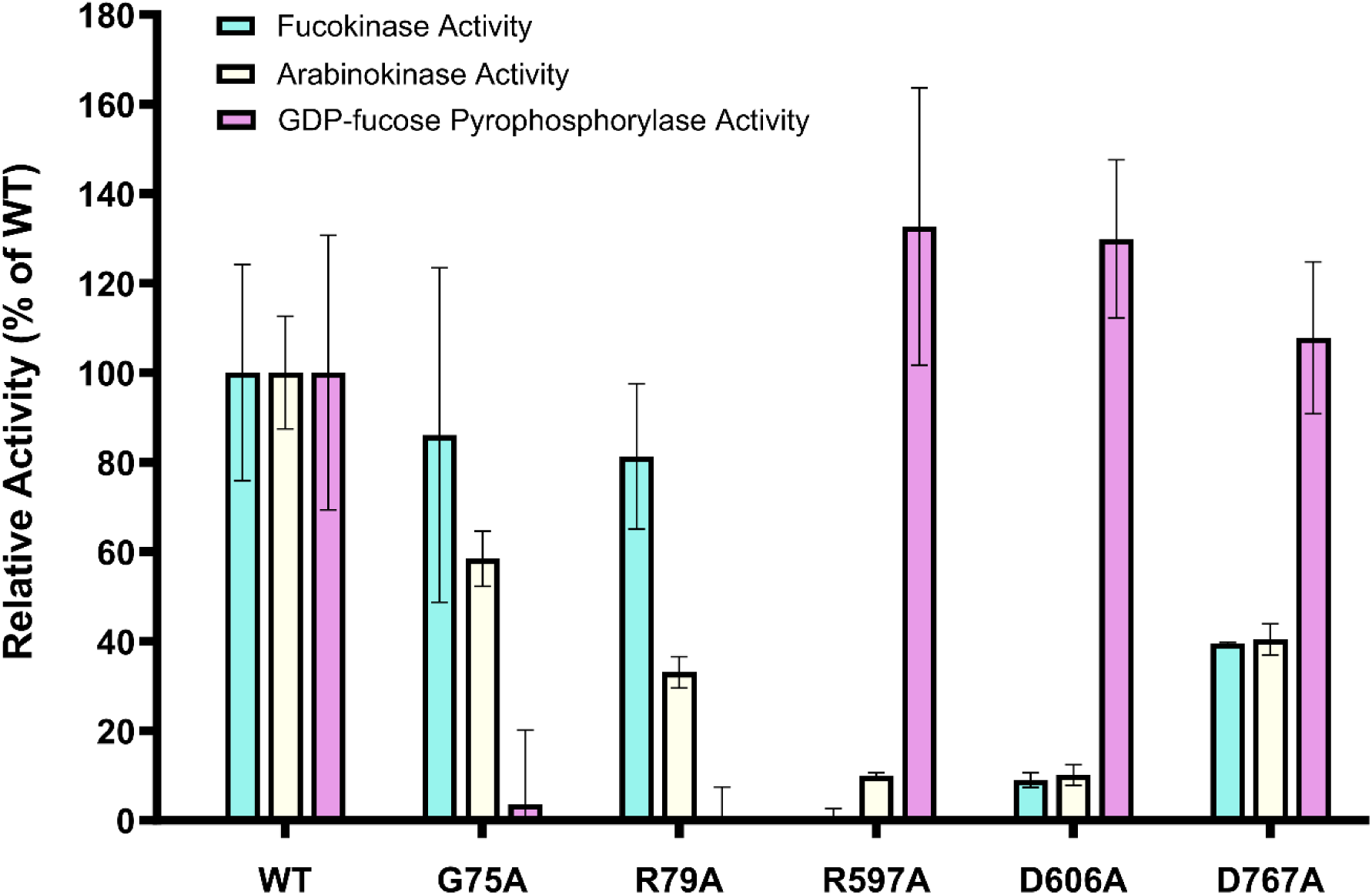
Relative activity of ***Tx***FKP mutants. The activity of each mutant relative to wildtype (WT) *Tx*FKP is shown. Each set of bars represents either the WT or mutant *Tx*FKP proteins. Bar heights represent the mean of at least three replicates conducted for fucokinase (cyan), arabinokinase (yellow), and pyrophosphorylase (pink) activity. The mean of each protein’s activity was normalized to the mean activity of WT *Tx*FKP, which is set to 100%. Standard deviations were calculated from the assay replicates and scaled proportionally to the normalized mean.

The most critical kinase residue identified in *Tx*FKP was R597, with the R597A-*Tx*FKP mutant losing all detectable fucokinase activity and nearly all arabinokinase activity. In the ColabFold2 model of *Tx*FKP, the side chain of this arginine protrudes deeply into the putative substrate-binding pocket that is also lined by D606 and D767 (Figure 2C). The analogous R592 residue in *Bf*FKP is positioned similarly and faces the phosphate group in the reported *Bf*FKP●Fuc-1-P complex (29). Taken together, this suggests that R597 is the critical phosphate-coordinating group that stabilizes the transition state of the phosphoryl transfer to the sugar within the active site of *Tx*FKP (42). It is unclear why the R597A- *Tx*FKP mutant retained some arabinokinase activity, but perhaps the lack of the methyl substituent at C5 permits D-Ara more freedom to sample different binding motifs wherein another residue can serve as a substitute for R597 and “rescue” the arabinokinase activity of an otherwise inactive mutant.

The *Tx*FKP variants with mutations in the *N*-terminus, G75A-*Tx*FKP and R79A-*Tx*FKP, lost nearly all pyrophosphorylase activity relative to the wildtype. The backbone amide of the analogous glycine (G76) in *Bf*FKP is known to form a hydrogen bond with the ribose of GTP, while the analogous arginine (R80) is predicted to facilitate the removal of the pyrophosphate moiety from GTP (29). Thus, the severe loss of activity in the G75A-*Tx*FKP and R79A-*Tx*FKP mutants suggests a similar critical role for these residues in *Tx*FKP. Additionally, the analogous glycine (G133) in the *Arabidopsis At*FKGP is critical for pyrophosphorylase activity (22), making this a conserved residue that FKPs from Clades A and B use to coordinate GTP.

Although some mutants of *Tx*FKP behaved as predicted, this was not the case with all of the tested variants. The D606A-*Tx*FKP mutant retained ∼10% of kinase activity, and the D767A-*Tx*FKP mutant retained ∼40% of kinase activity (Figure 4). The analogous D601A-*Bf*FKP mutant was reported to be catalytically inactive, while the analogous D762A-*Bf*FKP mutant retained only ∼5% of the fucokinase activity of the wildtype (28). Since D601A-*Bf*FKP has a reduced affinity for fucose but not ATP (29), we infer that the loss of activity in the D606A-*Tx*FKP mutant is likely attributed to impaired L-Fuc binding. In *Bf*FKP, D762 is predicted to coordinate L-Fuc and also act as the Brønsted base that deprotonates the C1 hydroxyl group of L-Fuc (29). However, since the D767A-*Tx*FKP mutant retained significant kinase activity, this residue cannot be critical for kinase activity in *Tx*FKP. These findings suggest that the *Tx*FKP aspartate residues D606 and D767 contribute primarily to L-Fuc binding, with a less significant role in the catalytic phosphoryl group transfer.

Testing the mutants for their ability to phosphorylate either L-Fuc or D-Ara also yielded a surprising result: mutations in the GFPP domain had a much more pronounced effect on arabinokinase activity than on fucokinase activity. The G75A-*Tx*FKP and R79A-*Tx*FKP mutants only exhibited ∼60% and ∼30% of the wildtype arabinokinase activity, despite maintaining at least 80% of the wildtype fucokinase activity (Figure 4). These mutations are distal to the kinase domain and their influence on kinase activity in a substrate-specific manner is intriguing. This result shows that FKP kinase specificity could be more intricately linked to the GFPP domain than previously appreciated. While further study is required to evaluate the regulatory “crosstalk” of the two domains of *Tx*FKP more systematically, one possible explanation is that these residues influence the conformational dynamics or the oligomeric states of the protein that permit *Tx*FKP to turn over D-Ara faster than L-Fuc (Table 1). As no other FKP mutants have been tested with both L-Fuc and D-Ara, it remains unclear whether these interdomain effects are conserved in other FKPs, or if this property has emerged only in *Tx*FKP, in conjunction with its faster turnover of D-Ara.

### Domain-specific FKP thermostability

Finally, thermal shift assays were used to determine how elevated temperature might have influenced the stability of *Tx*FKP compared to *Bf*FKP. Contrary to our predictions, we did not find that *Tx*FKP was a more thermostable protein than *Bf*FKP under standard conditions. Instead, at pH 7, both FKPs produced melt curves with biphasic peaks, seemingly corresponding to two unfolding events and two *T*m values per protein. For clarity, we refer to the lower *T*m as *T*m1 and the higher *T*m as *T*m2. For wildtype *Tx*FKP, *T*m1 corresponds to a slower unfolding than *T*m2, while the reverse occurs for *Bf*FKP, resulting in melt curve profiles resembling mirror images of one another (Figure 5). The *Tx*FKP melt curve also suggested another unfolding event at ∼68 °C, which was predicted to arise due to denaturation of the suspected GroEL contaminant. To verify this, we heat-shocked the sample at 50 °C for twelve hours, removed precipitated protein via centrifugation, and repeated the thermal shift assay using the soluble fraction. This heat-shocked sample contained only the contaminant protein and yielded a single *T*m of ∼68 °C (Figure S14), confirming that the signal observed at this temperature was indeed caused by the denaturation of the contaminant protein.

**Figure 5.**
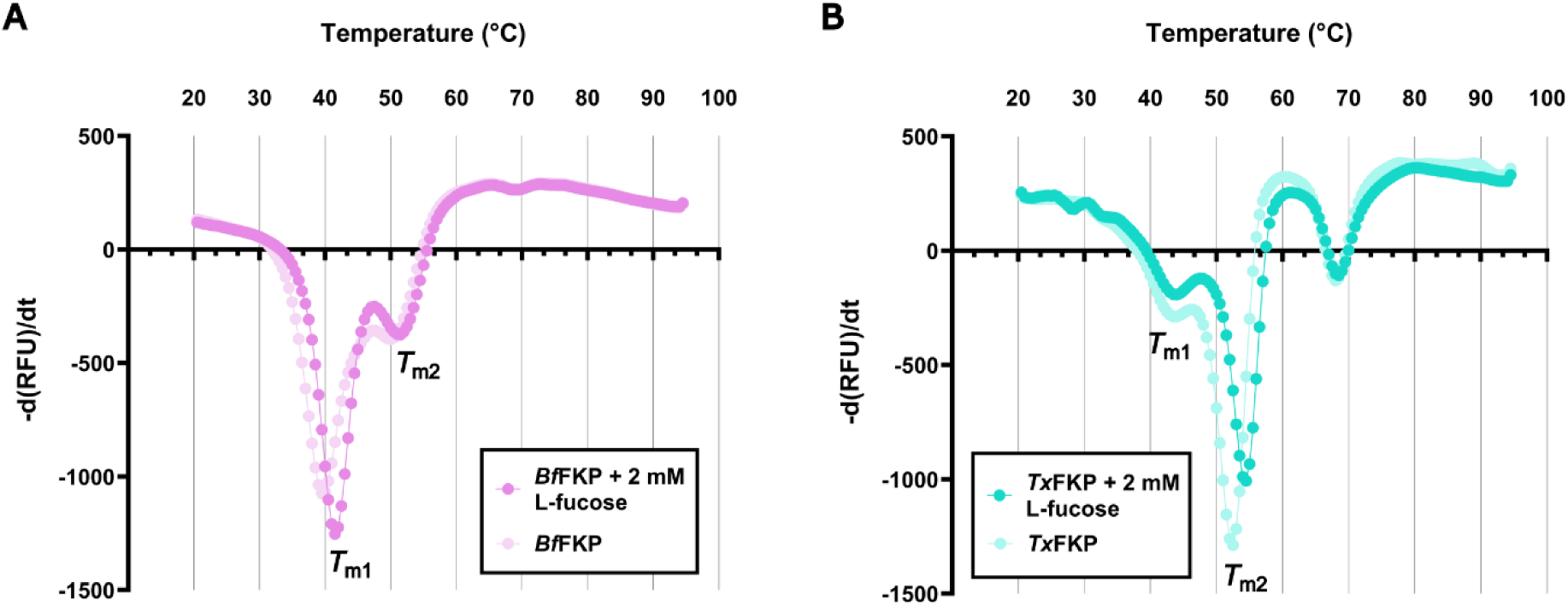
Thermal shift profiles of FKP proteins. The melting temperatures of *Bf*FKP and *Tx*FKP, with and without 2 mM L-fucose, are shown with the negative rate of change of SYPRO orange fluorescence as a function of temperature. Troughs represent a maximum rate of change, corresponding to the melting temperature (designated *T*m1 or *T*m2). Each point is the mean of at least three replicates. (A) Profiles for *Bf*FKP with (pink) and without (light pink) 2 mM L-Fuc. (B) Profiles for *Tx*FKP with (cyan) and without 2 mM L-Fuc (light cyan).

To investigate the possibility that each *T*m corresponded to an individual domain of the FKPs, we incubated the proteins with L-Fuc and repeated the thermal shift assay. Both FKP enzymes were stabilized by the addition of L-Fuc in a concentration-dependent manner, and this thermal shift is attributable to binding of these substrates. Incubation of *Tx*FKP with 2 mM L-Fuc did not affect *T*m1, but *T*m2 was increased by 2 °C (Table 2). None of the *Tx*FKP variants with point mutations in the kinase domain (R597A, D606A, D767A) demonstrated any thermal shift in response to incubation with L-Fuc.

**Table 2.**
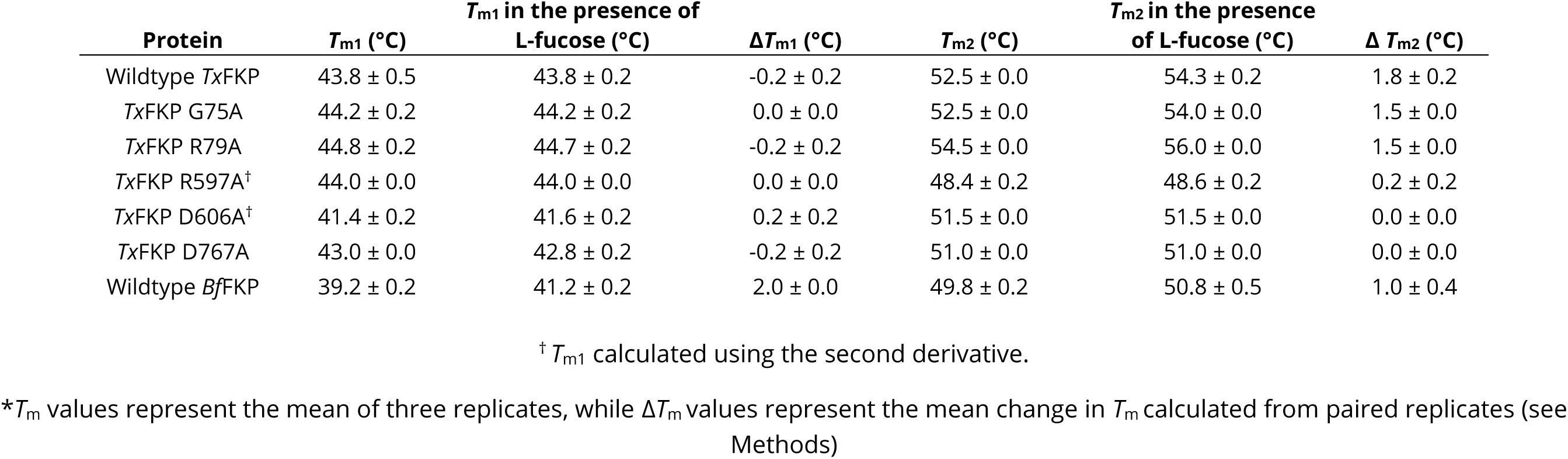
Average melting temperatures of FKP enzymes.

This indicates that each of these mutants has a reduced affinity for L-Fuc relative to the wildtype protein, providing additional evidence that these residues contribute to L-Fuc binding. *Tx*FKP variants with mutations in the pyrophosphorylase domain (G75A, R79A) responded similarly to the wildtype protein, exhibiting an increased *T*m2 in the presence of L-Fuc. Collectively, these results lead to the suggestion that *T*m2 corresponds to the FUK domain of *Tx*FKP: only *Tx*FKP mutants with perturbations in the FUK domain gave *T*m2 values that were unresponsive to L-Fuc incubation, and the FUK domain is the only expected L-Fuc binding location. By exclusion, *T*m1 then likely corresponds to the GFPP domain of the protein.

The response of *Bf*FKP melt curves to L-Fuc incubation was more complex: *T*m1 and *T*m2 of *Bf*FKP shifted by 2.0 °C and 1.0 °C, respectively, when incubated with L-Fuc (Table 2). While this prevented us from associating *T*m1 and *T*m2 with individual domains, this finding indicated that L-Fuc stabilized both domains of *Bf*FKP, suggesting a global conformational change in response to substrate binding. Such a conformational change would have the result of effectively “tightening” both domains, resulting in the observed shifts in both *T*m1 and *T*m2 of *Bf*FKP. Conformational changes are exploited by bifunctional enzymes like mammalian kinase/RNase Ire1 (43) and *Arabidopsis* lysine-ketoglutarate reductase/saccharopine dehydrogenase (44), and indeed conformational flexibility was recently hypothesized to assist substrate binding and release in *Bf*FKP (29). A recent report suggested that *Bf*FKP does not possess a substrate tunnel to transport Fuc-1-P from the FUK active site to the GFPP active site (29), so this conformational change may be a key feature in the *Bf*FKP-catalyzed salvage reaction.

The contrast between *Bf*FKP and *Tx*FKP responses to L-Fuc binding thus presents a potential difference between how these enzymes have evolved, specifically in how the proteins allow for an efficient transfer of Fuc-1-P from one domain to the other. It is possible that *Tx*FKP does not require a conformational change upon L-Fuc binding and has evolved a different strategy for this process. Other bifunctional enzymes such as formiminotransferase-cyclodeaminase (45) glutamine phosphoribosylpyrophosphate aminotransferase (46), tryptophan synthase (47), and thymidylate synthase/dihydrofolate reductase (48) facilitate ligand transport from one domain to another through hydrophobic tunnels or on the surface of a protein with electrostatic interactions. Further studies may confirm if *Tx*FKP employs a similar strategy, helping to explain if this feature evolved as an adaptation to higher temperatures.

## CONCLUSION

Our *in silico* analyses suggest that FKP enzymes originated from one common FKP ancestor and expand on their evolution by showing that FKP enzymes are absent from some organisms but frequently encoded by others. To determine how environmental factors might select for undiscovered traits in FKP enzymes, we characterized the second known prokaryotic FKP enzyme, *Tx*FKP, from the thermophile *T. xiamenensis*. Despite differences in host lifestyle, the kinetic parameters of *Tx*FKP kinase and pyrophosphorylase activities were similar to those of *Bf*FKP, and surprisingly, the two enzymes did not exhibit significantly different thermostabilities. However, *Tx*FKP demonstrated a modular, domain-specific thermostability and showed no evidence of a global conformational change upon L-Fuc binding. This result, along with the finding that mutations of proposed pyrophosphorylase catalytic residues affects the kinase activity differently with L-Fuc and D-Ara, provides new insight to the use of interdomain communication in the FKP family. Structural studies using different FKP enzymes may help uncover the evolutionary relevance of strategical differences in substrate transport between FKP domains. Additionally, the characterization of FKP proteins from different bacteria or even eukaryotes could reveal how conserved these properties are. Such insight would enhance current GDP-Fuc synthesis protocols and also address the question of why bifunctional FKP proteins appear to be selected for and maintained in some organisms.

## EXPERIMENTAL PROCEDURES

### General materials

Unless otherwise stated, *Escherichia coli* strains were grown at 37 °C on Lysogeny Broth (LB) agar with the following antibiotics as needed: chloramphenicol (Cm; 5 µg/mL), ampicillin (Amp; 100 µg/mL), kanamycin (Km; 25 µg/mL). Antibiotic concentrations were halved in liquid media.

### Phylogenetic analysis of FKP proteins

To gain insight into the evolutionary origins of bifunctional FKP proteins, we first downloaded and compiled Hidden Markov Models (HMM) from Pfam-A version 36.0 (49) and TIGRFAMs version 15.0 (50), as well as from the Cluster of Orthologous Genes (COGs) (51) and Protein Clusters (PRK) databases of NCBI’s Conserved Domain Database (CDD). The *Bf*FKP protein sequence was queried against all HMMs using the hmmscan function of HMMER version 3.3 (52), identifying PF07959 and COG2605 as the highest-scoring models to characterize GFPP and FUK domains, respectively.

PRK13412 was the highest-scoring model for full-length bifunctional FKPs.

Next, hmmsearch (52) was used to query the UniRef50 database for protein clusters matching the COG2605, PF07959, and PRK13412 models, and hmmscan was used to search the full HMM library to identify the top scoring HMMs for each cluster. To investigate the FUK domain, clusters that had COG2605 or PRK13412 as their top-scoring model, and clusters that had PF07959 and either COG2605 or PRK13412 in their top two scoring models, were extracted. To investigate the GFPP domain, clusters that had PF07959 or PRK13412 as their top-scoring model, and clusters that had COG2605 and either PF07959 or PRK13412 in their top two scoring models, were extracted. Putative FKP proteins were defined based on the presence of all three of the desired HMMs in their top three highest-scoring models. Sequences below 333 residues (for FUKs), 406 residues (for GFPPs), and 739 residues (for full-length FKPs), and sequences above 1745 residues (for all datasets), in length were removed. The filtered set of sequences were aligned using MAFFT version 7.471 (53), alignments were trimmed using trimAl version 1.4.rev22 with the automated1 option (54), and the trimmed alignments were tested against several substitution models (LG, WAG, JTT, Q.pfam, JTTDCMut, DCMut, VT, PMB, Blosum62, Dayhoff) with IQ-TREE 2 version 2.2.2.4 (32, 55). For each dataset, IQ-TREE2 was used to generate a maximum-likelihood phylogeny with the best model and 1000 ultrafast bootstrap replicates (32). As the bi-domain nature of FKP proteins confounds the selection of an outgroup, we also used IQ-TREE2’s non-reversible amino acid model NONREV to predict rootsrap support values for each dataset. The topology of the final phylogenetic trees was based on where the rootstrap value was highest (73% for the FKP phylogeny and 96% for the FUK phylogeny), except for in the GFPP phylogeny, which received ambiguous support and was rooted based on the FKP monophyletic clade or at its midpoint. Phylogenetic trees were visualized using iTOL version 6 (56).

### FKP structural predictions and comparison

ColabFold2 version 1.5.5 (57) was used to predict the monomeric structures of *Bf*FKP and *Tx*FKP. The highest-ranked structures were aligned using the matchmaker feature in ChimeraX version 1.6.1 (58) using the default settings and *Bf*FKP as the reference protein.

### Sequence similarity networks

Protein sequences, as acquired by the methods listed above, were analyzed using the all-by-all BLAST function of the Enzyme Function Initiative-Enzyme Similarity Tool (EFI-EST) server (59, 60). Sequence similarity networks (SSNs) were then generated from this dataset using an alignment score of 100 for the FKP dataset, 70 for the FUK dataset, and 100 for the GFPP dataset, corresponding to respective sequence identities of 30%, 40%, and 35%. SSNs were visualized using Cytoscape version 3.9.1 (61).

### Plasmid construction and site-directed mutagenesis

Genes encoding *Bf*FKP and *Tx*FKP were codon-optimized for expression in *E. coli*, then synthesized within pET-28a expression vectors by Bio Basic (Table S1). The pET-28a construct containing the *fkp* gene from *T. xiamenensis* was used as a template for inverse polymerase chain reaction (PCR). Primers (Table S2) designed to induce point mutations in the GFPP domain (G75A, R79A) and in the FUK domain (R597A, D606A, D767A) were synthesized with 5’ phosphate groups by Thermo Fisher. PCR amplicons were digested with DpnI (New England Biolabs) to remove methylated DNA, circularized using T4 ligase (New England Biolabs), and transformed into *E. coli* DH10B cells. Successfully mutated plasmids were identified using whole plasmid Oxford Nanopore Technologies sequencing at Plasmidsaurus (Louisville, KY, USA).

### Recombinant protein expression and purification

Competent *E. coli* BL21 (DE3) cells were co-transformed with each respective pET plasmid and a pACYC vector encoding the chaperone proteins GroEL, GroES, and Trigger Factor (TF) (62) (Table S1). One-litre liquid cultures of each strain were grown until they reached an optical density at 600 nm (OD600) of 0.4-0.8, then expression was induced with 1 mM IPTG. The recombinant proteins were expressed at 16 °C and shaken at 180 RPM for a duration of approximately 20 hours.

All following centrifugation and chromatography was performed at 4 °C. Induced cultures were harvested by centrifugation at 4,347 x *g* for ten minutes. Cell pellets were resuspended in lysis buffer (20 mM Tris-HCl, 100 mM NaCl, 10 % glycerol (v/v), pH 7.6) and 1 mg/mL lysozyme (BioShop). Samples were mechanically lysed with three passes through an Emulsiflex C5 (Avestin) or a French Press (Sigma Aldrich), then clarified by centrifugation at 16,000 x *g* for one hour. A Bio-Rad Econo-Column was packed with 5 mL Ni sepharose resin (Cytiva) and pre-equilibrated with lysis buffer before adding cell lysate and slowly shaking the mixture for five minutes. Three 30 mL volumes of wash buffer (20 mM Tris-HCl, 100 mM NaCl, 20 mM imidazole, 10 % glycerol (v/v), pH 7.6) were run through the column, and the nickel-bound protein was then eluted in elution buffer (20 mM Tris-HCl, 100 mM NaCl, 500 mM imidazole, 10 % glycerol (v/v), pH 7.6). The eluted samples were concentrated using Amicon Ultra centrifugal filter units (Sigma). Finally, buffer exchange and de-salting were performed using PD-

10 columns (Cytiva) according to the manufacturer’s protocol. Purified protein was eluted in storage buffer (40 mM Tris-HCl, 100 mM NaCl, 10 % glycerol (v/v), pH 7.6) and flash frozen in liquid nitrogen. The total protein concentrations were measured using the Pierce bicinchoninic acid (BCA) Assay Kit (Thermo Fisher). For the calculation of FKP present, the intensity of Coomassie Brilliant Blue G-250 staining on an SDS-PAGE gel was measured using ImageJ version 1.54h (63).

### HILIC-HPLC-MS Product Detection

To characterize the products of the *Tx*FKP-catalyzed reaction(s), reactions were based on a recent report (6). L-Fuc or Fuc-1-P were dissolved in HEPES buffer (100 mM, pH 8.0) to a final concentration of 10 mM. To this mixture, 15 mM ATP, 10 mM GTP, and various combinations of 10 mM MgCl2, 10 mM MnCl2, or 5 mM EDTA were added. Both ATP and GTP were included in the two-step reactions, while only GTP was included to test the pyrophosphorylase activity alone. The inorganic pyrophosphatase from *Pasteurella multocida Pm*PpA (40), purified as described previously (64), was added at a concentration of 5 µg per µmol of starting sugar to promote the forward reaction. *Bf*FKP or *Tx*FKP were added at 1 mg/ml, then reactions were incubated at 37 °C for one hour, quenched by centrifugation with a 10 kDa MWCO filter (VWR), and analyzed using hydrophilic interaction liquid chromatography mass spectrometry (HILIC-HPLC-MS) (37, 38).

Analytical HILIC-HPLC-MS was performed on an Agilent InfinityLab LC/MSD series with a Waters XBridge BEH Amide column, 5 μm, 4.6 x 250 mm at a flow rate of 2.0 mL/min, injection volume of 20 μL (2 mg/mL), with 1% of the flow diverted to the ESI-MS detector using a splitter. Mobile phase A was 10 mM ammonium formate in water, adjusted to pH 6.5 with formic acid; mobile phase B was 10 mM ammonium formate in 90% acetonitrile (pH = 7.4). The linear gradient used is detailed in Table S3.

### Kinase kinetic assays

To monitor the FUK-catalyzed reaction, FUK activity was coupled to the activities of pyruvate kinase (PK) type II (Sigma-Aldrich) and lactate dehydrogenase (LDH) type II (Sigma-Aldrich), largely as described previously (26). Reactions consisted of 10 mM MnCl2, 1 mM phosphoenolpyruvate (PEP), 300 µM NADH, 25 U/mL PK, 35 U/mL LDH, and varied concentrations of ATP, L-Fuc, and D-Ara, in reaction buffer (50 mM Tris, 100 mM NaCl, pH 7.5). To determine the affinity for L-fuc or D-Ara, the ATP concentration was held at 5 mM for the *Tx*FKP reactions and 3 mM for the *Bf*FKP reactions (to avoid the observed inhibitory effect). To characterize affinity for ATP, the L-Fuc concentration was held at 1 mM for *Tx*FKP and *Bf*FKP reactions. When evaluating the activity of *Tx*FKP mutants, sugars were used at concentrations >5x the *K*M values exhibited by the wildtype enzyme and ATP was used at 5 mM (*Tx*FKP) or 3 mM (*Bf*FKP).

Reaction mixtures were pre-incubated at 37 °C before initiating reactions with the addition of 277 nM FKP. Using an Epoch2 plate reader (BioTek), the absorbance at 340 nm was measured every ten to fifteen seconds for several minutes at 37 °C. Absorbance values were converted to concentrations using an NADH standard curve.

### Pyrophosphorylase kinetic assays

The GFPP-catalyzed reaction was monitored using a modified phosphate detection assay (41). The inorganic pyrophosphatase *Pm*PpA (40) was used to hydrolyze the inorganic pyrophosphate (PPi) produced during the GFPP-catalyzed reaction, resulting in an inorganic phosphate (Pi) signal that was detected using a malachite green reagent (Cell Signaling Technology). Reactions consisted of 10 mM MgCl2, 0.05 mg/mL *Pm*PpA, 40 nM of FKP, and varied concentrations of Fuc-1-P (Biosynth) and GTP (Sigma Aldrich), in reaction buffer (50 mM Tris, 100 mM NaCl, pH 7.5). The concentration of *Pm*PpA was sufficiently high as to not limit the rate of the GFPP-catalyzed reaction. To characterize affinity for GTP, Fuc-1-P concentration was held at 250 µM, while to characterize affinity for Fuc-1-P, GTP concentration was held at 100 µM for TxFKP and 400 uM for *Bf*FKP. When evaluating the activity of *Tx*FKP mutants, the Fuc-1-P and GTP concentrations were 250 µM and 200 µM, respectively.

Reactions were initiated by the addition of either Fuc-1-P or GTP and incubated at 37 °C. Three aliquots of 25 µL were removed at 15-30 second intervals during the linear phase of the reactions (approximately one minute) and added to 100 µL of malachite green reagent. Colour development was allowed to proceed for five or fifteen minutes before reading the absorbance at 635 nm in an Epoch2 plate reader (BioTek). Absorbance values were converted to concentrations using a Pi standard curve specific to the incubation time.

### Thermal shift assays

To investigate differences in the melting temperature (*T*m) between enzymes, they were incubated with SYPRO orange (Bio-Rad) (65), and thermally denatured. Mixtures comprised of 400 µg/mL FKP enzyme, 5 X SYPRO orange (Thermo Fisher), and buffer (50 mM Tris, 50 mM NaCl, pH 7.0) were loaded into a PCR plate (Bio-Rad). When investigating stabilizing effects of co-incubation with the substrate, mixtures included 2 mM L-Fuc. Samples were allowed to equilibrate to room temperature for five minutes, then the plate was sealed and heated from 20 °C to 95 °C in a qPCR thermal cycler (Bio-Rad). Fluorescence measurements were taken during ten-second 0.5 °C intervals. Data were collected using Bio-Rad’s CFX Maestro software. To calculate melting temperatures, the negative first or second derivative of SYPRO orange fluorescence was plotted against temperature. For each protein, samples were measured in triplicate in the presence and absence of L-Fuc. Standard deviations were calculated for the mean *T*m, and for the mean change in *T*m (Δ*T*m) of the paired replicates.

## DATA AVAILABILITY

All scripts necessary to repeat the phylogenetic analyses in this study are publicly available on GitHub at https://github.com/diCenzo-Lab/013_2025_Fkp_analyses. Interactive versions of all phylogenetic trees presented here can be accessed through iTOL at https://itol.embl.de/shared/1tzSBFA310S3y.

## SUPPORTING INFORMATION

This article contains supporting information (28, 53, 62, 66, 67).

