## Supplemental Information for "Characterization of bacterial fucokinase/GDP-fucose pyrophosphorylase (FKP) enzymes supports the evolution of interdomain communication and modularity in the FKP family"

**This file includes:**

Tables S1 to S4 (pages S-2 to S-5)
Figures S1 to S14 (pages S-6 to S-19)
References for the Supporting Information (page S-20)

### SUPPORTING TABLES

**Table S1. Plasmids and bacterial strains used in this study**

| Plasmid/Strain Name | Characteristics | Reference |
|---|---|---|
| ***Escherichia coli*** | | |
| BL21 (DE3) | *fhuA2 [lon] ompT gal* (*λ DE3*) *[dcm] ΔhsdS* | New England Biolabs |
| DH10β | *Δ(ara-leu) 7697 araD139 fhuA ΔlacX74 galK16 galE15 e14- ϕ80dlacZΔM15 recA1 relA1 endA1 nupG rpsL (Str$^R$) rph spoT1 Δ(mrr-hsdRMS-mcrBC)* | New England Biolabs |
| B006 | BL21 (DE3) (pGQ0097,P003); Amp$^R$ Cm$^R$ | This work |
| B009 | BL21 (DE3) (pGQ0097,pGQ0167); Km$^R$ Cm$^R$ | This work |
| EcGQ0250 | BL21 (DE3) (pGQ0097,pGQ0169); Km$^R$ Cm$^R$ | This work |
| EcGQ0251 | BL21 (DE3) (pGQ0097, pGQ0173); Km$^R$ Cm$^R$ | This work |
| EcGQ0252 | BL21 (DE3) (pGQ0097,pGQ0172); Km$^R$ Cm$^R$ | This work |
| EcGQ0253 | BL21 (DE3) (pGQ0097, pGQ0171); Km$^R$ Cm$^R$ | This work |
| EcGQ0254 | BL21 (DE3) (pGQ0097,pGQ0168); Km$^R$ Cm$^R$ | This work |
| **Plasmids** | | |
| P003 | pET-15b vector containing *BfFKP* gene; Amp$^R$ | Yi et al (1) |
| pGQ0097 | pACYC vector encoding GroEL-EsTF chaperone genes; Cm$^R$ | Lammpa et al (2) |
| pGQ0167 | pET-28a vector containing *TxFKP* gene; Km$^R$ | This work |
| pGQ0168 | pGQ0167 containing *TxFKP* gene with G75A mutation; Km$^R$ | This work |
| pGQ0169 | pGQ0167 containing *TxFKP* gene with R79A mutation; Km$^R$ | This work |
| pGQ0171 | pGQ0167 containing *TxFKP* gene with R597A mutation; Km$^R$ | This work |
| pGQ0172 | pGQ0167 containing *TxFKP* gene with D606A mutation; Km$^R$ | This work |
| pGQ0173 | pGQ0167 containing *TxFKP* gene with D767A mutation; Km$^R$ | This work |

Amp – ampicillin; Cm – chloramphenicol; Km – kanamycin

**Table S2. Oligonucleotide primers used in this study**

| Primer Name | Primer Sequence (5'-3') | Usage |
|---|---|---|
| AH001 | GCG**CA**CAGTCCCGCCGTTTAC | Forward primer to induce G75A mutation in pET28a-*Tx*FKP plasmid |
| AH002 | CGGCGTGAATAATGATCTTTTTGTCCGC | Reverse primer to induce G75A mutation in pET28a-*Tx*FKP plasmid |
| AH003 | CCTACGCGCCCCTTGGGA | Forward primer to induce R79A mutant in pET28a-TxFKP plasmid |
| AH004 | CTGGTAAA**GC**GCGGGACTGG | Reverse primer to induce R79A mutant in pET28a-TxFKP plasmid |
| AH007 | GCC**GCA**TTGGACCTGGCAG | Forward primer to induce R597A mutant in pET28a-TxFKP plasmid |
| AH008 | CGGACTACGTCCCCAGACG | Reverse primer to induce R597A mutant in pET28a-TxFKP plasmid |
| AH009 | CG**C**CACGCCACCTTACTGC | Forward primer to induce D606A mutation in pET28a-*Tx*FKP plasmid |
| AH010 | GACCAACCTCCTGCCAGGTCC | Reverse primer to induce D606A mutation in pET28a-*Tx*FKP plasmid |
| AH011 | GGATGGCAGG**C**CCAGTACGGAGG | Forward primer to induce D767A mutation in pET28a-*Tx*FKP plasmid |
| AH012 | GCCGCCAGTGGTAAGAAGCTGTTCC | Reverse primer to induce D767A mutation in pET28a-*Tx*FKP plasmid |

*Bolded and underlined nucleotides indicate regions not homologous to the template. Primers were synthesized with 5' phosphate groups.

**Table S3. Linear gradient conditions for HILIC-HPLC-MS**

| Time (min) | A (%) | B (%) |
|---|---|---|
| 0 | 0 | 100 |
| 10 | 0 | 100 |
| 20 | 10 | 90 |
| 40 | 70 | 30 |
| 50 | 90 | 10 |
| 60 | 0 | 100 |

*Mobile phase A was 10 mM ammonium formate in water, adjusted to pH 6.5 with formic acid; mobile phase B was 90% acetonitrile with 10 mM ammonium formate in water (pH = 7.4)

**Table S4. Recombinant protein expression yields**

| Protein | Total Yield (mg/L culture) | %FKP: %GroEL | FKP-Weighted Yield (mg/L culture) |
|---|---|---|---|
| *Bf*FKP | 20.9 | 96:4 | 20.1 |
| *Tx*FKP | 14.4 | 55:45 | 7.9 |
| G75A-*Tx*FKP | 13.5 | 52:48 | 7.0 |
| R79A-*Tx*FKP | 14.5 | 52:48 | 7.5 |
| K88A-*Tx*FKP† | N.D. | N.D. | N.D. |
| R597A-*Tx*FKP | 8.0 | 60:40 | 4.8 |
| D606A-*Tx*FKP | 10.7 | 59:41 | 6.3 |
| D767A-*Tx*FKP | 11.9 | 52:48 | 6.2 |

†K88A-*Tx*FKP did not express at a level sufficient for purification

### SUPPORTING FIGURES

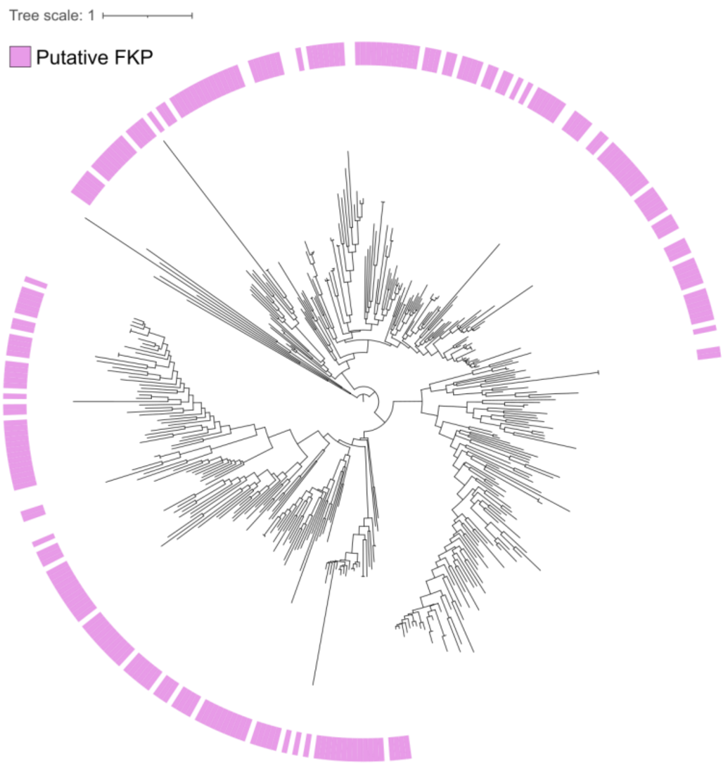


**Figure S1. GFPP phylogeny with midpoint root.** A maximum-likelihood phylogeny generated from UniRef50 hits to the GFPP protein family. The scale bar represents the average number of amino acid substitutions per site, node support was calculated based on 1000 ultrafast bootstrap replicates, and an interactive form with bootstrap supports can be accessed at itol.embl.de/shared/1tzSBFA310S3y. The phylogeny is identical to the one reported in Figure 1B, but it is rooted at its midpoint. Protein clusters that also have predicted FUK domains, and thus are putative FKP proteins, are indicated with a pink rectangle.

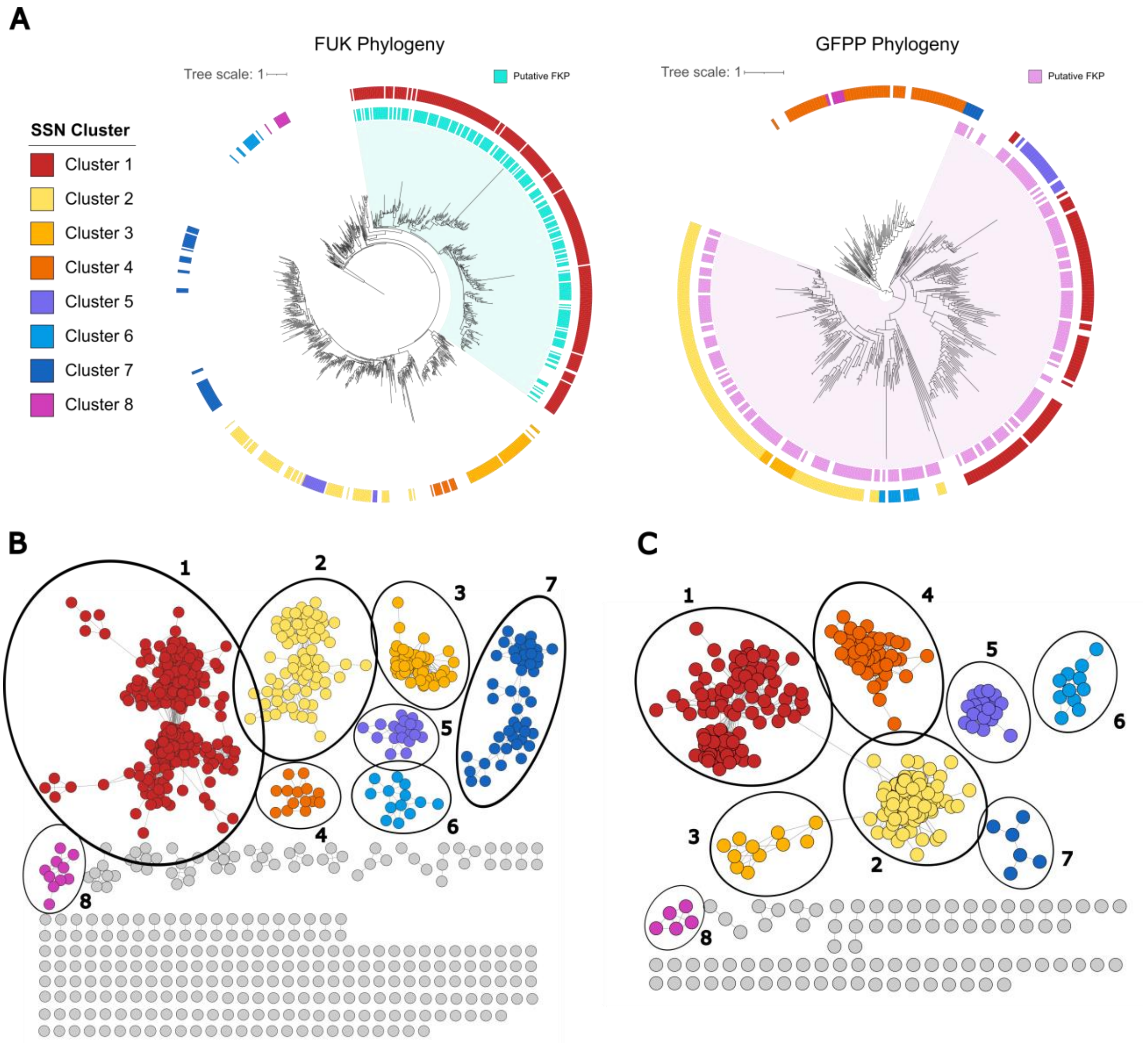


**Figure S2. FUK and GFPP phylogenies with sequence similarity clustering.** Phylogenies and accompanying sequence similarity networks (SSNs) for FUK and GFPP proteins are shown. The phylogenies are identical to those reported in Figure 1. For all phylogenies the scale bar represents the average number of amino acid substitutions per site, node support was calculated based on 1000 ultrafast bootstrap replicates, and interactive forms with bootstrap supports can be accessed at itol.embl.de/shared/1tzSBFA310S3y. (**A**) Maximum-likelihood phylogenies for FUK (left) and GFPP (right) proteins. Putative FKP proteins are indicated with a cyan or magenta rectangle, and the “FKP monophyletic clade” is highlighted in pale cyan or pale magenta. The multicoloured data strip indicates membership to a cluster from the corresponding SSN. (**B**) SSN for the FUK protein set. Clusters of eight UniRef50 accessions or less are not coloured. Nodes (circles) represent UniRef50 accessions, and the connecting edges (lines) represent a shared identity of approximately 40% or above. (**C**) SSN for the GFPP protein set. Clusters of three UniRef50 accessions or three or less are not coloured. Nodes (circles) represent UniRef50 accessions, and the connecting edges (lines) represent a shared identity of approximately 35% or above.

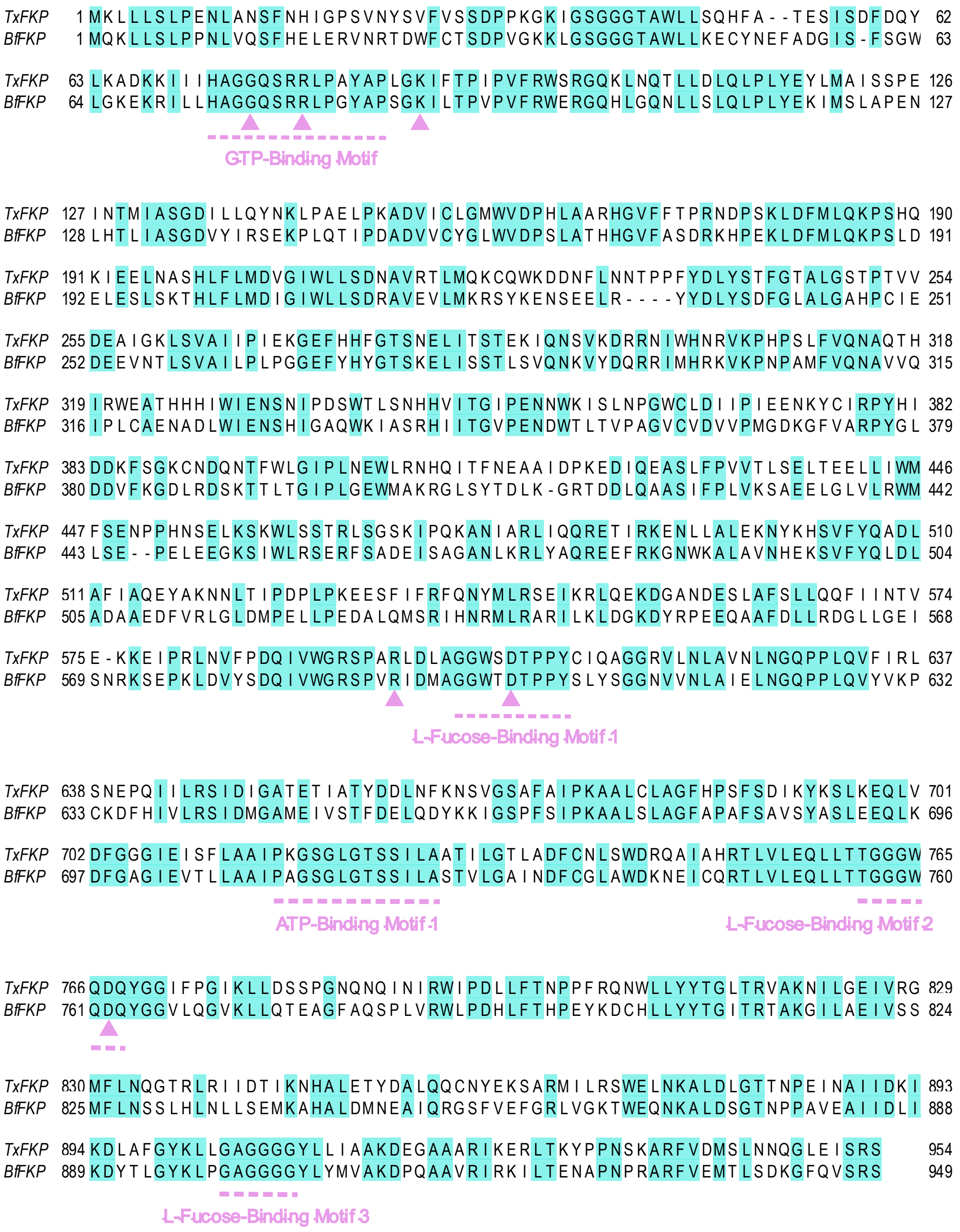


**Figure S3. Sequence alignment of *Bf*FKP and *Tx*FKP protein sequences**. *Bf*FKP and *Tx*FKP amino acid sequences were aligned using MAFFT version 7.471 (3) and visualized with Jalview version 2.11.4.0 (4). Residues that are conserved between the two proteins are highlighted in cyan. Substrate-binding motifs identified by Liu et al (2019) (5) are indicated with a dashed magenta line. Magenta arrows designate residues that are important for either GFPP or FUK activity in *Bf*FKP (5), and which were selected for mutagenesis in *Tx*FKP in this study.

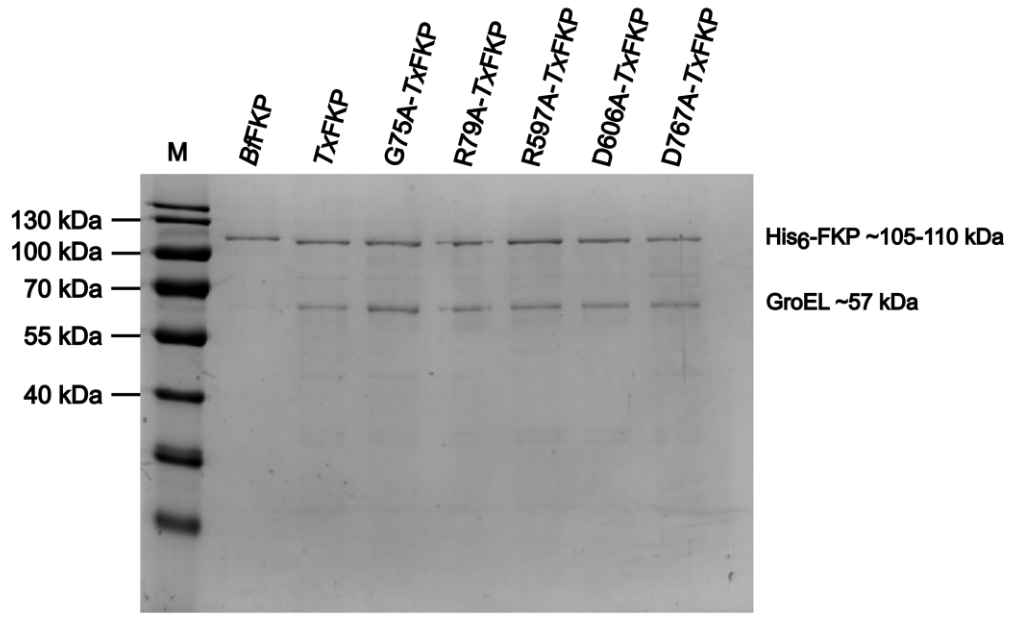


**Figure S4. SDS-PAGE gel of FKP proteins.** A 10% SDS-PAGE gel loaded with 100 ng of each recombinantly expressed and purified FKP protein and stained with Coomassie Brilliant Blue G-250. An **M** is used to indicate the PageRuler Prestained Protein Ladder (Thermo Fisher). Expected masses of FKP proteins and the GroEL chaperone are indicated on the right.

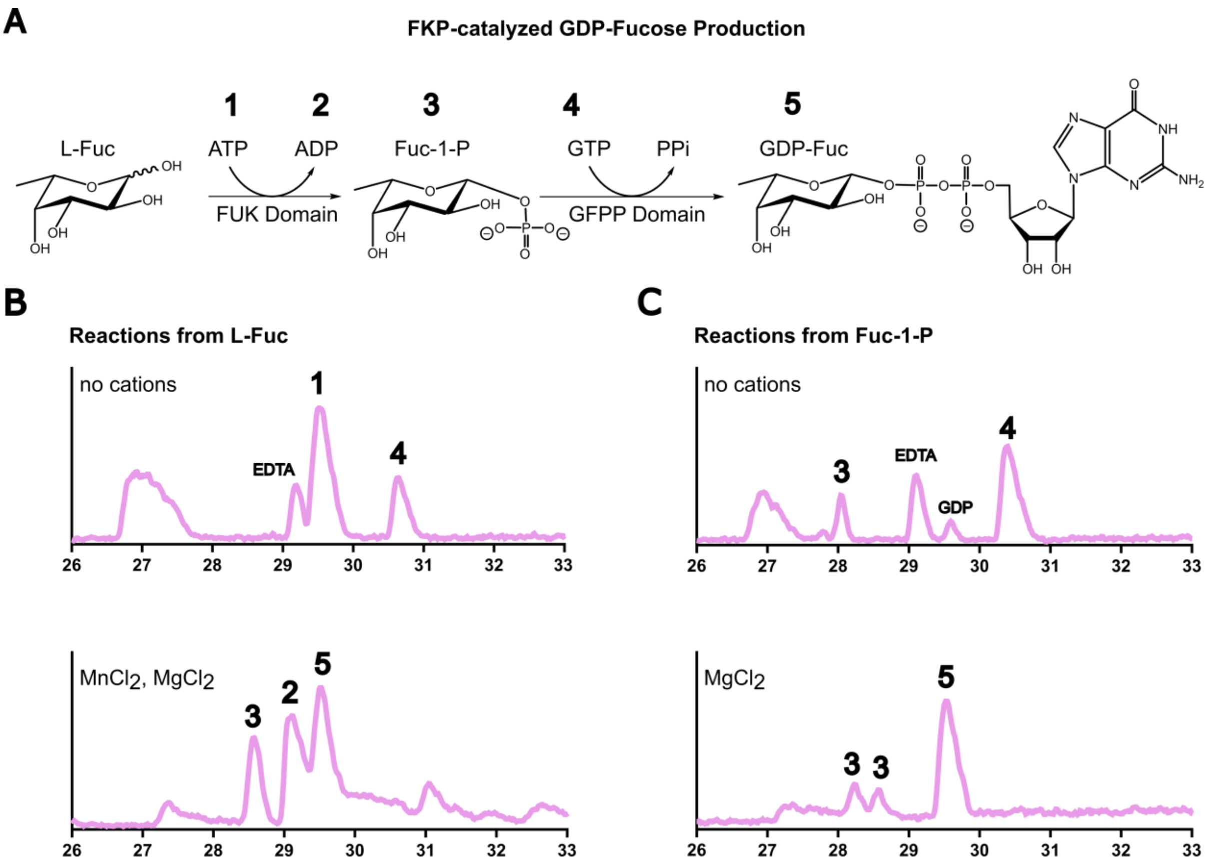


**Figure S5. *Bf*FKP-catalyzed GDP-fucose synthesis.** HILIC-HPLC results demonstrating the *Bf*FKP-catalyzed synthesis of GDP-Fuc are shown. Numbers from one to five represent a reaction substrate or product as follows: (1) ATP, (2) ADP, (3) Fuc-1-P, (4) GTP, and (5) GDP-Fuc. Mass spectrometry chromatograms (negative mode), with signal intensity plotted against time, are shown of reaction mixtures following one hour of incubation at 37 °C. All chromatograms were standardized based on the retention time of HEPES (23-25 min). The salt(s) used in each condition are indicated on the chromatograms, and the “no cations” condition included 5 mM EDTA. Relevant peaks are numbered accordingly and the peak corresponding to EDTA is labelled where applicable. (**A**) A reaction scheme representing the FUK and GFPP-catalyzed reactions during the two-step synthesis of GDP-Fuc. (**B**) Chromatograms of reaction mixtures where *Bf*FKP was incubated with L-Fuc as a starting substrate. (**C**) Chromatograms of reaction mixtures where *Bf*FKP was incubated with Fuc-1-P as a starting substrate.

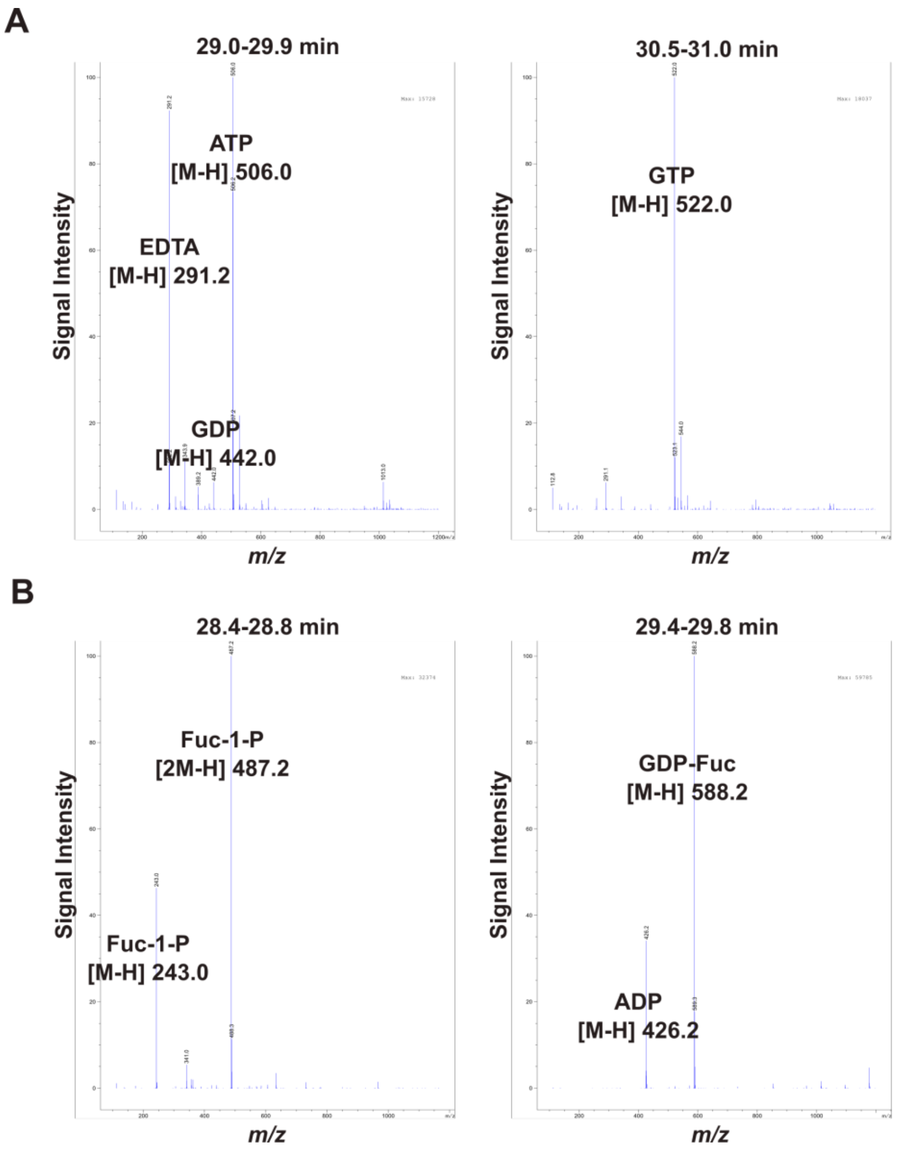


**Figure S6. Mass spectrometry characterization of *Bf*FKP reaction products with L-Fuc.** *Bf*FKP reaction mixtures using L-Fuc as the starting sugar were incubated for one hour, then the reaction products were analyzed with HILIC-HPLC-MS. Mass spectrometry was performed using the negative ion mode. Each panel shows the total ion chromatogram (TIC) corresponding to the time frame when the relevant compounds eluted. All retention times are standardized based on the retention of HEPES. The panels A and B refer to reactions using different cation conditions as follows: (**A**) 5 mM EDTA was included, and (**B**) 10 mM of both $MnCl_2$ and $MgCl_2$ were included.

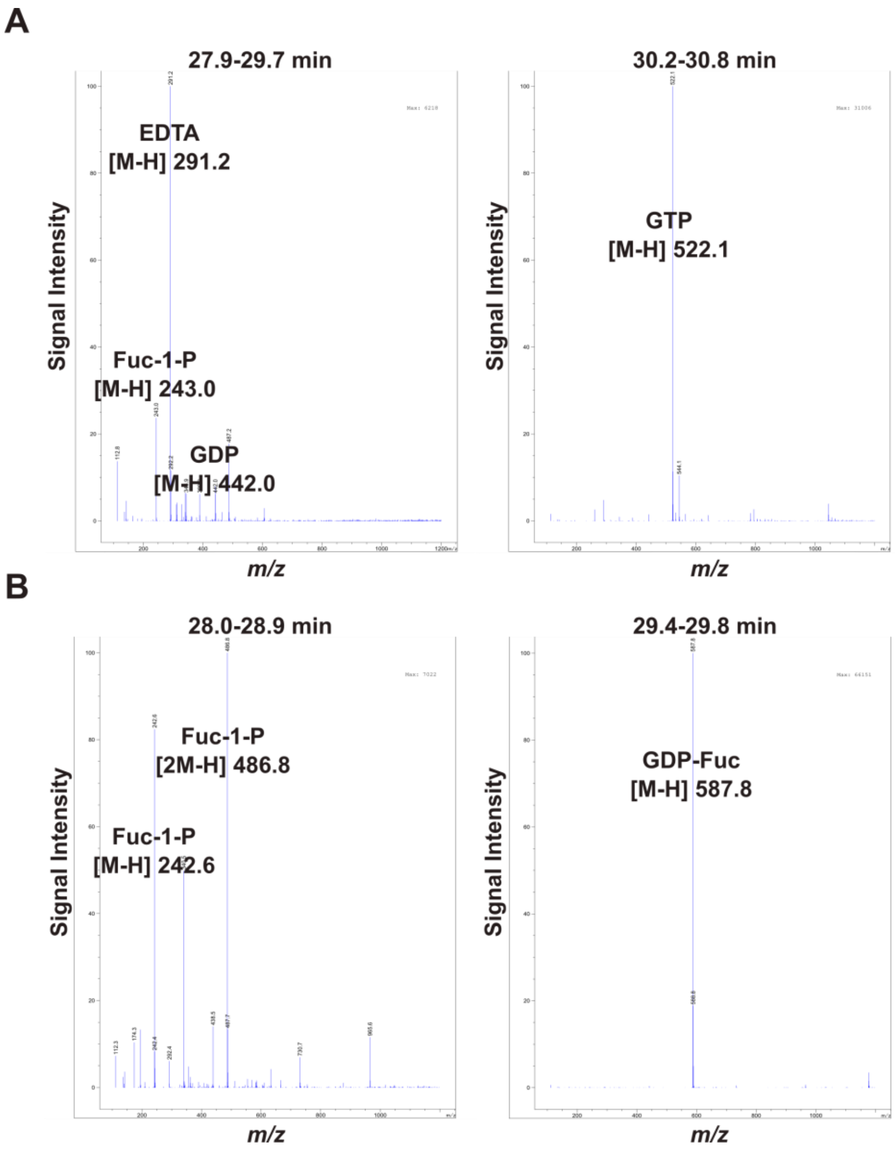


**Figure S7. Mass spectrometry characterization of *Bf*FKP reaction products with Fuc-1-P.** *Bf*FKP reaction mixtures using Fuc-1-P as the starting sugar were incubated for one hour, then the reaction products were analyzed with HILIC-HPLC-MS. Mass spectrometry was performed using the negative ion mode. Each panel shows the total ion chromatogram (TIC) corresponding to the time frame when the relevant compounds eluted. All retention times are standardized based on the retention of HEPES. The panels A and B refer to reactions using different cation conditions as follows: (**A**) 5 mM EDTA was included, and (**B**) 10 mM of $MgCl_2$ was included.

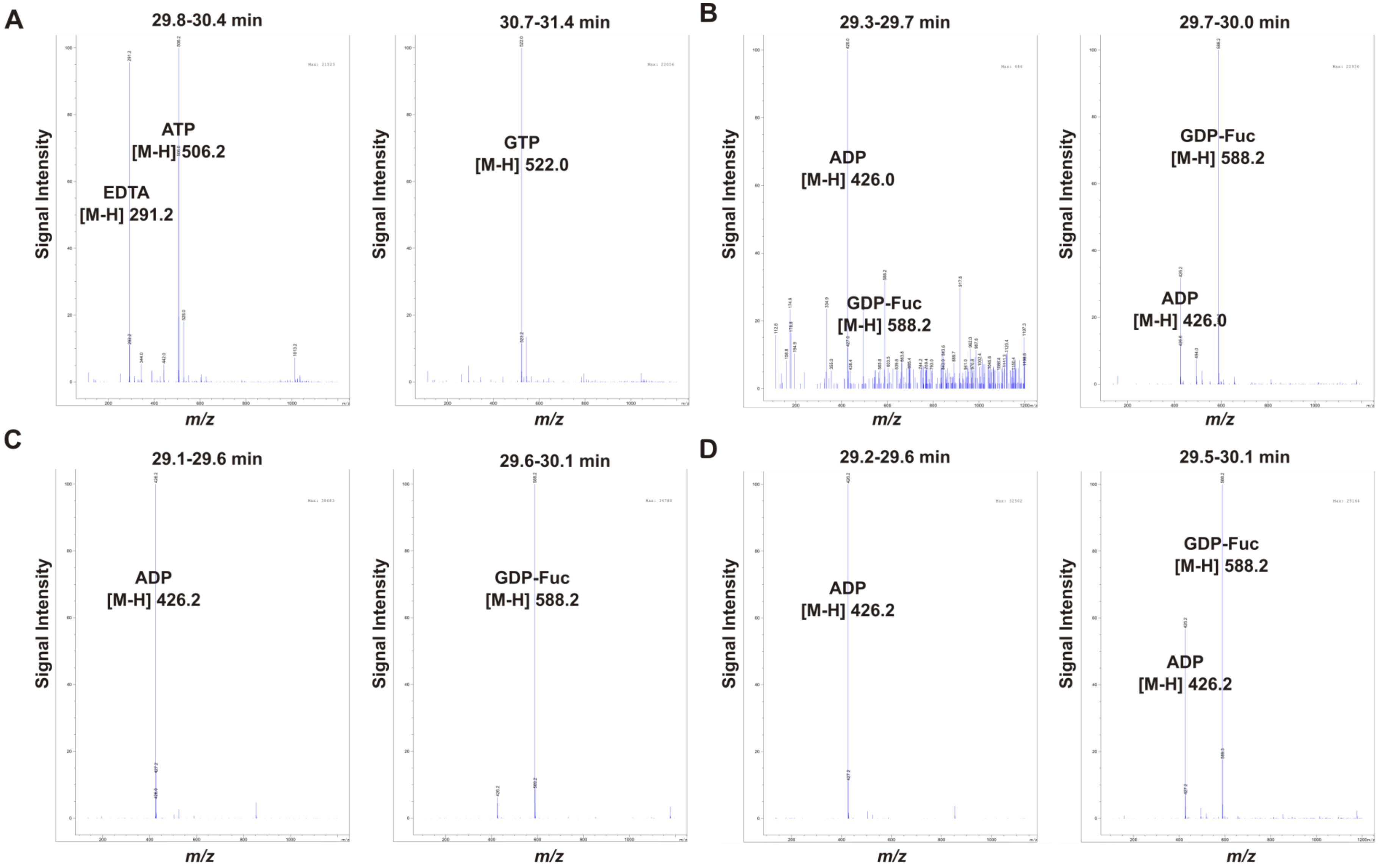


**Figure S8. Mass spectrometry characterization of *Tx*FKP reaction products with L-Fuc.** *Tx*FKP reaction mixtures using L-Fuc as the starting sugar were incubated for one hour, then the reaction products were analyzed with HILIC-HPLC-MS. Mass spectrometry was performed using the negative ion mode. Each panel shows the total ion chromatogram (TIC) corresponding to the time frame when the relevant compounds eluted. All retention times are standardized based on the retention of HEPES. The panels A, B, C, and D refer to reactions using different cation conditions as follows: (**A**) 5 mM EDTA was included, (**B**) 10 mM of both $MnCl_2$ and $MgCl_2$ were included, (**C**) 10 mM of $MnCl_2$ was included, and (**D**) 10 mM of $MgCl_2$ was included.

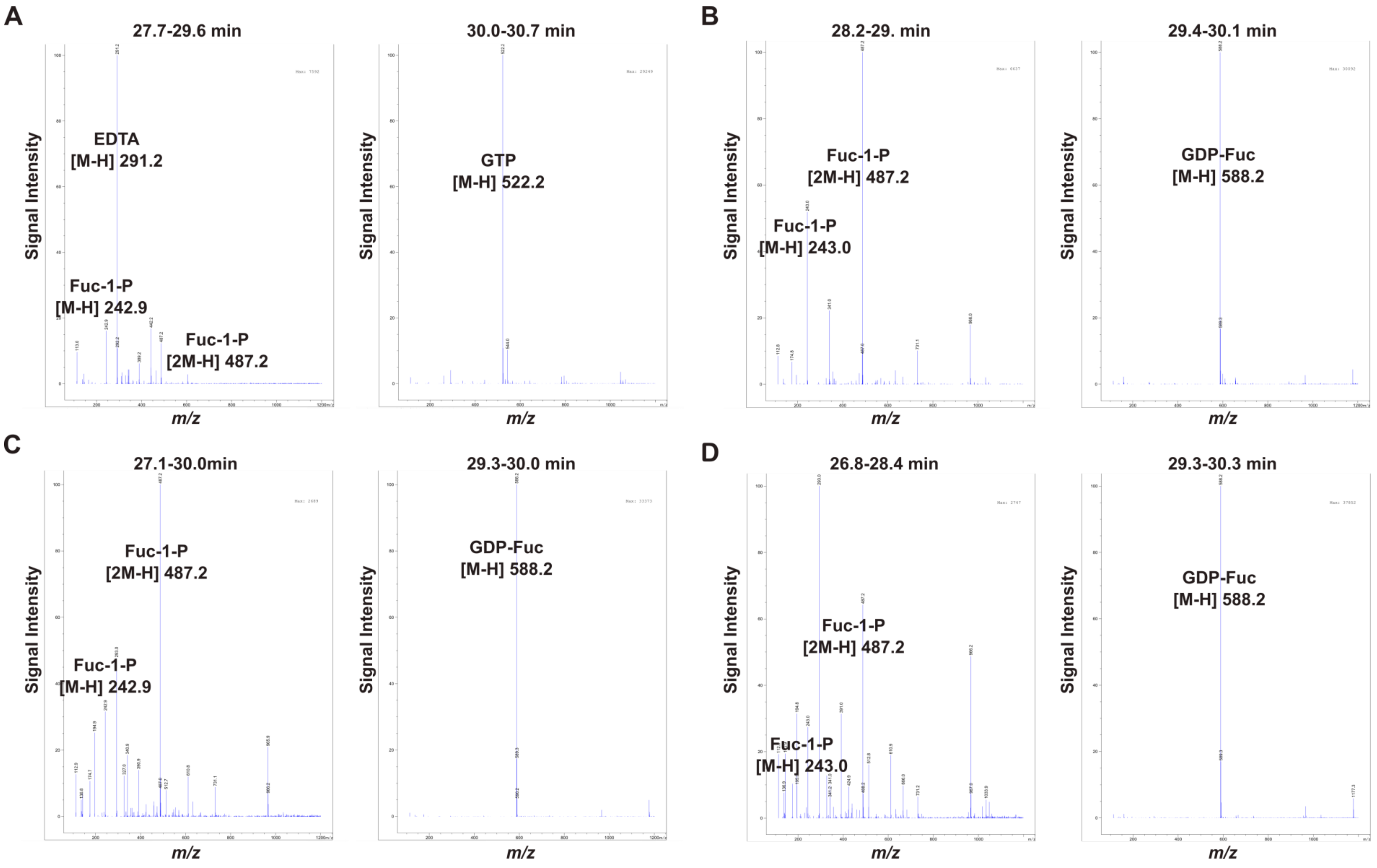


**Figure S9. Mass spectrometry characterization of *Tx*FKP reaction products with Fuc-1-P.** *Tx*FKP reaction mixtures using Fuc-1-P as the starting sugar were incubated for one hour and the reaction products were analyzed with HILIC-HPLC-MS. Mass spectrometry was performed using the negative ion mode. Each panel shows the total ion chromatograms (TICs) corresponding to the time frames when the relevant compounds eluted. All retention times are standardized based on the retention of HEPES. The panels A, B, C, and D refer to reactions using different cation conditions as follows: (**A**) 5 mM EDTA was included, (**B**) 10 mM of both $MnCl_2$ and $MgCl_2$ were included, (**C**) 10 mM of $MnCl_2$ was included, and (**D**) 10 mM of $MgCl_2$ was included**.**

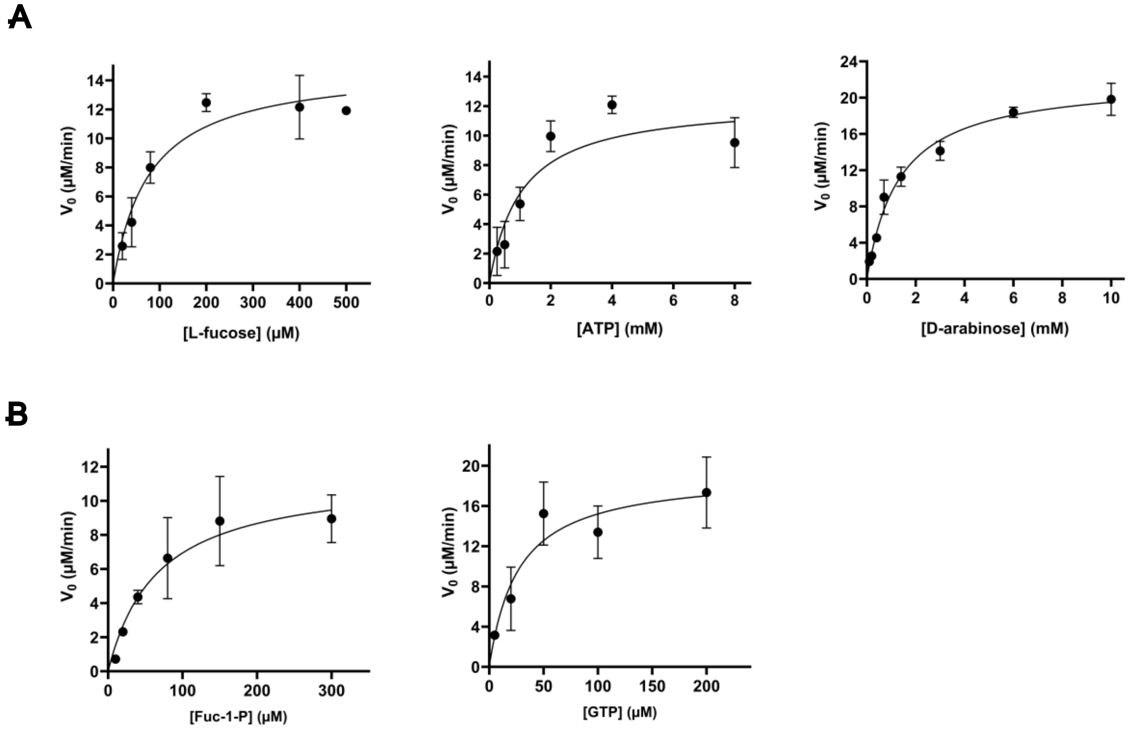


**Figure S10. Michaelis-Menten plots of *Tx*FKP activity.** The initial reaction velocities are plotted against substrate concentration for each indicated substrate. Data were fitted to the Michaelis-Menten equation using the Least-Squares option on GraphPad Prism version 10.0.3. Each point represents the mean of at least three replicates, and error bars represent standard deviation from the mean. (**A**) Plots for the *Tx*FKP-catalyzed kinase activity. (**B**) Plots for the *Tx*FKP-catalyzed pyrophosphorylase activity.

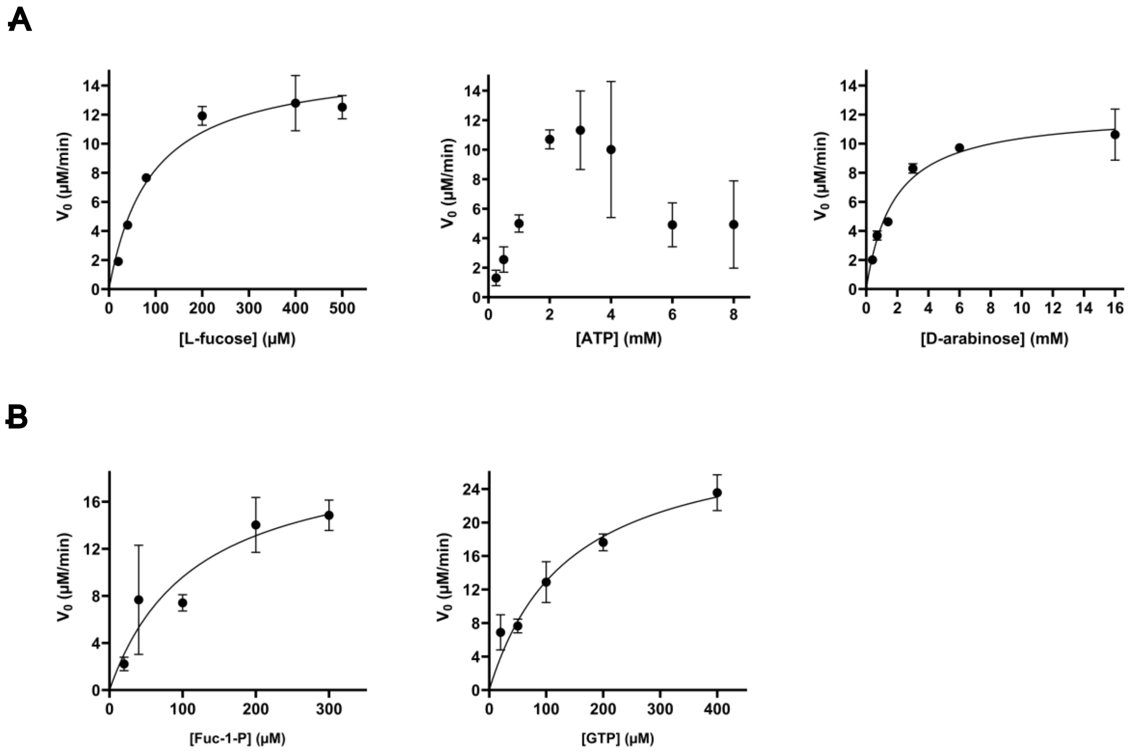


**Figure S11. Michaelis-Menten plots of *Bf*FKP activity.** The initial reaction velocities are plotted against substrate concentration for each indicated substrate. Data were fitted to the Michaelis-Menten equation using the Least-Squares option on GraphPad Prism version 10.0.3, except for the plot with varied ATP concentrations. Each point represents the mean of at least three replicates, and error bars represent standard deviation from the mean. (**A**) Plots for the *Bf*FKP-catalyzed kinase activity. (**B**) Plots for the *Bf*FKP-catalyzed pyrophosphorylase activity.

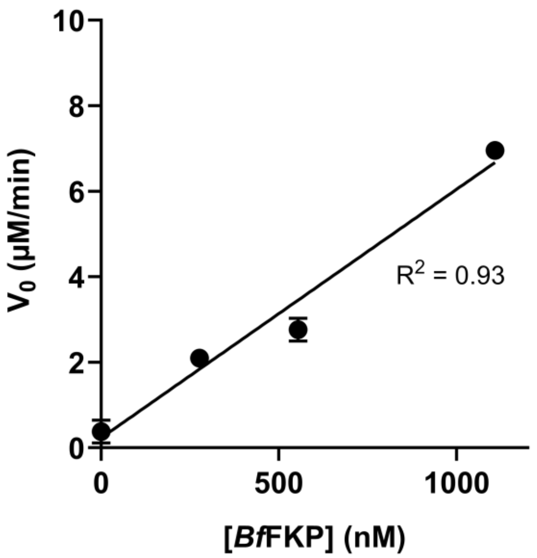


**Figure S12. [*Bf*FKP]-dependent FUK activity at 5 mM [ATP].** The initial fucokinase reaction velocity is plotted against the concentration of *Bf*FKP, when [ATP] was 5 mM and [L-fucose] was 1 mM. Data were fitted to a linear regression using the Least-Squares option on GraphPad Prism version 10.0.3. Each point represents the mean of three replicates, and the error bars represent standard deviation from the mean.

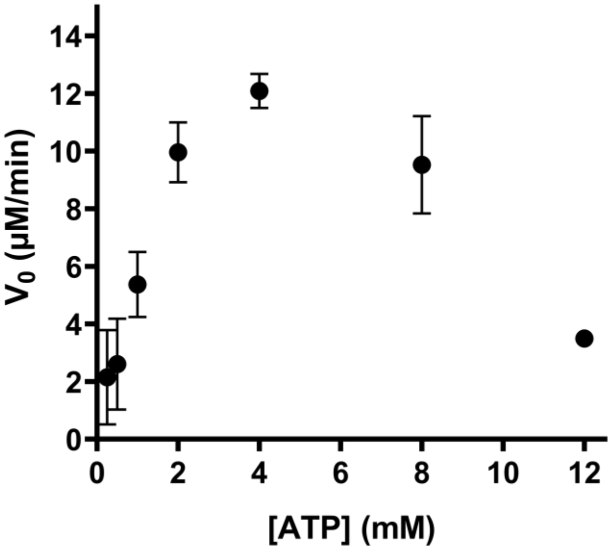


**Figure S13. *Tx*FKP FUK inhibition at high [ATP].** The initial velocity is plotted against ATP concentration. Each point represents the mean of at least three replicates, and error bars represent standard deviation from the mean.

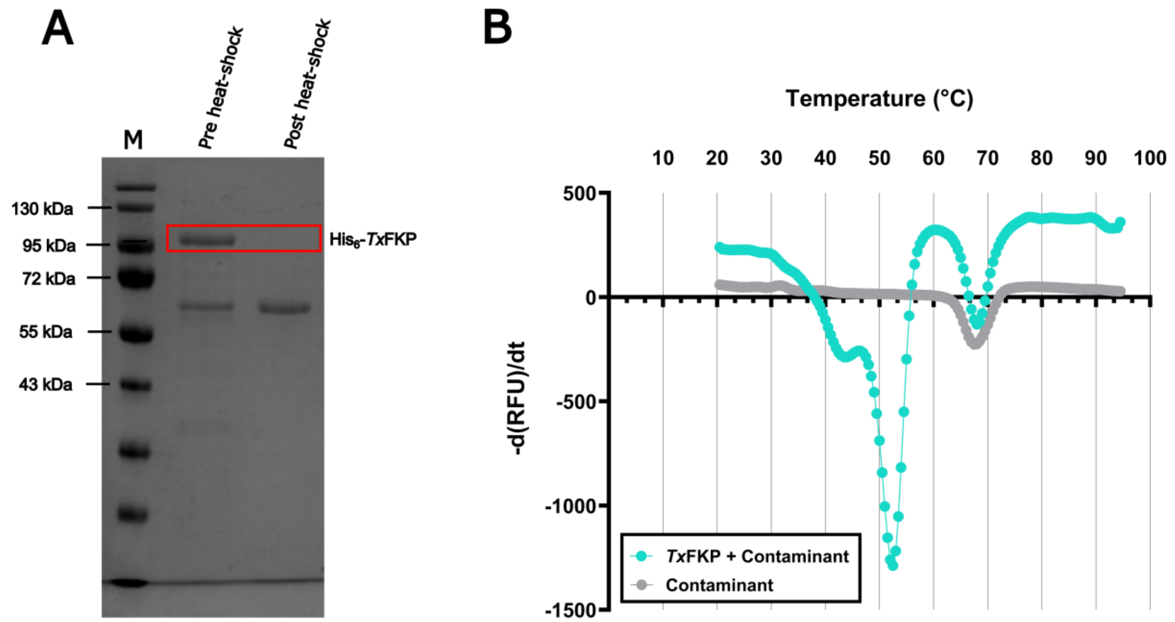


**Figure S14. Contaminant protein melting temperature**. (**A**) A 12% SDS-PAGE gel showing the removal of *Tx*FKP from the contaminant. From left to right, the lanes contain **M** (the PageRuler Prestained Protein Ladder), purified *Tx*FKP protein sample, and a sample following a twelve-hour 50 °C heat shock. (**B**) Melting temperatures of the pre-heat shock sample is shown in cyan, while the post-heat shock sample is shown in grey. The rate of change of SYPRO orange fluorescence is plotted against temperature. Points represent the average of at least three replicates.
